# Joint QTL mapping reveals patterns of pleiotropy across structured environments in budding yeast

**DOI:** 10.64898/2026.10.30.763211

**Authors:** Shreyas Gopalakrishnan, Eliot F. Fenton, Artur Rego-Costa, Samantha Petti, Michael M. Desai

**Affiliations:** Department of Molecular and Cellular Biology, Harvard University, Cambridge, MA 02138; Department of Physics, Harvard University, Cambridge, MA 02138; Department of Organismic and Evolutionary Biology, Harvard University, Cambridge, MA 02138; Department of Mathematics, Tufts University, Medford, MA 02155

**Keywords:** genotype-phenotype maps, complex traits, pleiotropy

## Abstract

Quantitative trait locus (QTL) mapping studies in model organisms have long been used to infer the structure of genotype-phenotype maps. However, prior work has largely focused on analyzing the genetic architecture of complex traits individually. Here, we sought to instead infer genotype-phenotype maps for a large set of related phenotypes simultaneously. To do so, we used bulk assays to measure the competitive fitness of each of 100,000 individuals from a large budding yeast QTL mapping panel across a set of 101 phenotypes related by a “hub-and-perturbation” pattern, and then used a forward search approach to jointly infer the genetic basis of all phenotypes together. Our results reveal complex patterns of pleiotropy in the genetic architecture of these traits. We find that these patterns of pleiotropy reflect the structure of the environmental relationships as well as functional information about individual variants and the interactions between them. This suggests that high-throughput QTL mapping across structured environments can help uncover biologically meaningful genetic interactions, along with their variation across related environmental contexts.

---

Quantitative trait locus (QTL) mapping studies have long been used to map the genetic architecture of complex traits in model organisms. Numerous researchers have used this approach in budding yeast to characterize genotypephenotype maps underlying many physiologically relevant traits (1–5). The basic logic of these studies is straightforward: we construct a mapping panel consisting of offspring derived from some mating scheme, assay genotypes and phenotypes of these offspring, and then use statistical associations between them to infer the structure of the genotype-phenotype map (6, 7). This work has shed important light on specific genes and pathways relevant for these traits, as well as the overall structure of genetic networks in this organism (8–10).

Our ability to draw insight from these mapping studies is limited by several practical experimental choices which constrain our ability to identify statistically significant associations between genotypes and phenotypes. This has motivated numerous recent studies aiming to improve the power and resolution of these studies (4, 11, 12). For example, advanced crosses can be used to increase the density of recombination breakpoints in a mapping panel, reducing linkage disequilibrium and hence enabling finer mapping of the causal variants underlying variation in phenotype (13). Other work has aimed to survey genetic variation more broadly by constructing large mapping panels from circular (14) or multiparental (15) crosses. More recent work by Nguyen Ba *et al*. (16) focused instead on increasing power and resolution simply by increasing sample size. These authors constructed a barcoded panel of about 100,000 haploid F1 budding yeast offspring of a cross between a wine strain and laboratory strain, sequenced each segregant, and measured a set of 18 growth rate phenotypes. The increased power afforded by the large sample size in this study allowed the authors to map the effects of a large number of small-effect QTLs, and to map pairwise epistatic effects.

These earlier studies have all provided important insights into the genetic basis of many important complex traits, and more generally into the structure of genetic networks in yeast and the functional consequences of natural variation. However, this work primarily focuses on analyzing each measured phenotype individually. In this study, we leverage the large mapping panel constructed by Nguyen Ba *et al*. (16) to instead *jointly* map the genetic basis of more than 100 related phenotypes. We chose these phenotypes to represent a “hub- and-perturbation” structure, involving varying the intensity of a set of specific perturbations (e.g. temperature) across each of several unrelated base media environments.

Our results provide to our knowledge the largest-scale data set yet available characterizing the pleiotropic effects of natural variants in budding yeast across a large set of more than 100 phenotypes. We then aim to use the observed correlations in genetic architecture across these phenotypes as the basis for characterizing structure in the organization of genetic networks in yeast. The essential idea is that QTLs whose effect sizes vary across phenotypes (e.g. across a temperature gradient) in similar ways are likely to be involved in the same or related biological pathways (e.g. relating to temperature response). This is similar in spirit to several recent studies that aim to identify lower-dimensional latent structure in the pleiotropic effects of specific mutations or QTLs (17, 18), and to previous work in yeast genetics using similarities in the patterns of epistatic interactions between mutations observed in large-scale gene deletion surveys to infer hierarchical modular organization in genetic networks (19, 20).

The organization of this paper is as follows. We begin in the next section by describing the large barcoded mapping panel that forms the basis for this study, the selection of the phenotypes we chose to measure, and our approach to measuring these phenotypes using bulk barcode-based relative growth rate assays. We then describe our approach to jointly inferring the genetic basis of these traits using a greedy forwardsearch algorithm. We next describe general properties of the QTLs underlying the genetic architecture of these complex traits. Finally, we analyze how the pleiotropic effects of these QTLs across related phenotypes follow characteristic patterns, and describe how these patterns may reflect aspects of the structure of genetic networks in yeast.

## A broad, structured survey of phenotype space

To survey pleiotropic structure across a large set of complex traits in *Saccharomyces cerevisiae*, we measured relative growth rate phenotypes (i.e. competitive fitness) of a large set of yeast strains (the barcoded mapping panel described below) across a set of 101 media conditions chosen to span a “hub-and-perturbation” pattern.

### A barcoded mapping panel

Our experiments here use a panel of *∼*100,000 haploid F1 offspring (“segregants”) generated from a cross between a standard laboratory yeast strain, BY4741 (henceforth BY), and a European wine strain, RM11-1a (henceforth RM) (21), created in earlier work by Nguyen Ba *et al*. (16). Each segregant is labeled with a unique 32-bp DNA barcode at a neutral locus. The basic cross design is summarized in Fig. 1A; see Materials and Methods in Nguyen Ba *et al*. (16) for more details. We note that the two parental strains differ at 41,594 single-nucleotide polymorphisms (SNPs) and show large differences in many phenotypes of interest; crosses between these parents have been extensively studied in previous work (4, 22, 23).

**Fig. 1.**
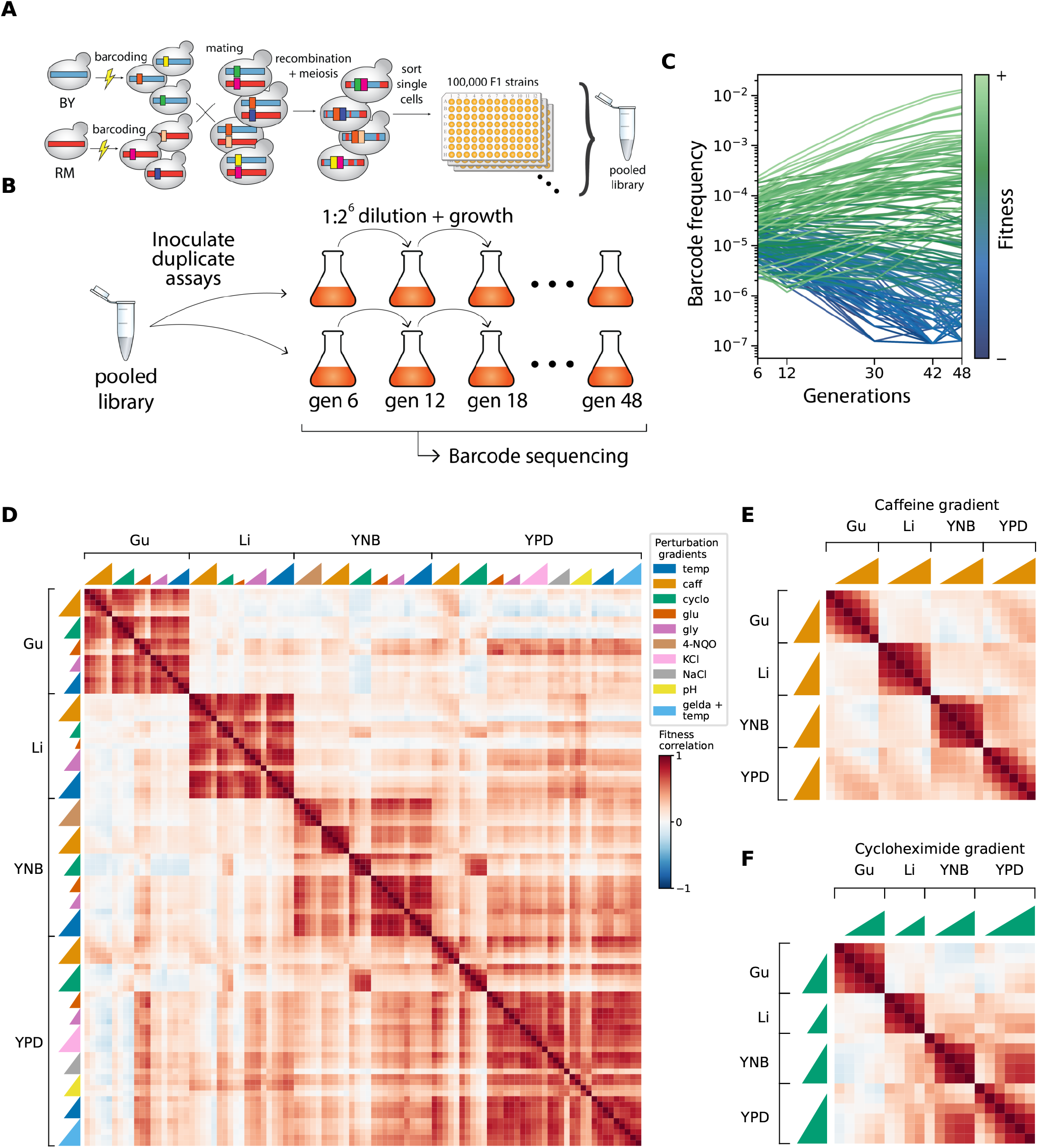
Experimental design and observed phenotypic correlations. **(A)** Schematic of the yeast cross design. Two diverged haploid yeast strains, BY4741 and RM11-1a, were transformed with diverse DNA barcodes at a neutral locus. The resulting pools of barcoded haploids were mated to produce heterozygous diploids, barcodes were moved onto a single chromosome using a Cre-Lox system, and diploids were then sporulated to produce a set of 100,000 uniquely barcoded haploid F1 offspring (segregants). Each segregant was whole-genome sequenced, and the association between segregants and barcodes was also identified. See (16) for more details. **(B)** Bulk fitness assays. For each environment, we inoculated the pooled library containing all 100,000 segregants into a flask containing 100 mL of the corresponding liquid medium and propagated in batch culture with daily 1:64 dilutions for eight days. We sampled an aliquot daily and deeply sequenced the barcode locus at five different timepoints to estimate the frequency of each barcoded segregant lineage over time. **(C)** Fitness inference. We infer the relative fitness of each segregant in that environment from the changes in each barcode frequency over time. **(D)** Observed correlations in fitness across all 101 environments. Each environment is identified by its hub, perturbation, and perturbation level. Details of these environments are listed in Table 1 and Table S1. **(E)** Heatmap of correlation in fitness across the caffeine perturbation gradient in all four hubs. **(F)** Heatmap of correlation in fitness across the cycloheximide perturbation gradient in all four hubs.

Each of the segregants in our panel has an average of *∼*50 recombination breakpoints, which means there are on average *∼*820 SNPs (or *∼*235 kb) between recombination breakpoints. Thus, the F1 genotypes contain long stretches of each parental genome, punctuated by the occasional breakpoint. This linkage structure was leveraged by Nguyen Ba *et al*. (16) to cheaply and efficiently genotype each segregant using lowcoverage sequencing (median coverage of *∼*0.6x). A separate amplicon-based sequencing approach was used to sequence the barcode region and to associate each barcode with the corresponding segregant genotype (16).

**Table 1.**
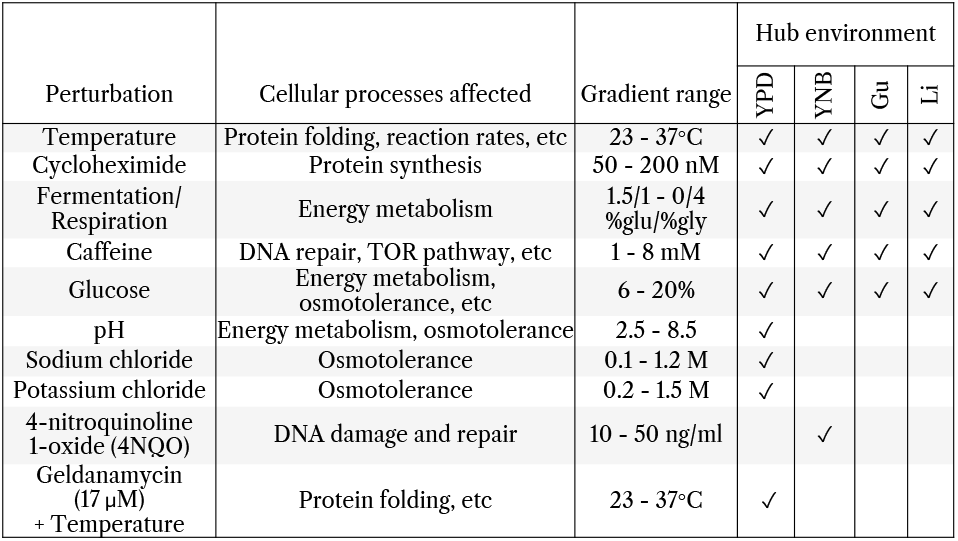
Summary of the 101 environments assayed in this study. The four hub environments are YPD (1% yeast extract, 2% peptone, 2% glucose), YNB (yeast nitrogen base without amino acids, with ammonium sulfate, and 2% glucose), Gu (YPD with 6 mM guanidinium chloride), and Li (YPD with 20 mM lithium acetate), all at 30^*°*^C. Each perturbation is added to the base medium of the indicated hub(s) at several levels spanning the gradient range shown. See Table S1 for the complete list of environments and media.

### Choice of media environments

We measured relative growth rate phenotypes for all *∼*100,000 segregants in each of 101 different media environments. Our choice of conditions was guided by earlier work that measured the competitive fitness of this panel of segregants in 18 different environments (16). This work found that this competitive fitness was highly correlated across 12 of these 18 environments (*R*^2^ *>* 0.5), suggesting that the differences between these conditions represent subtle perturbations to the genotype-to-phenotype map. Fitness in the other 6 environments was relatively uncorrelated both with each other and with the block of 12 correlated environments, suggesting that the differences between these conditions represent large shifts in the underlying genotype-phenotype map.

To probe the structure of pleiotropy across both subtle perturbations and large shifts in conditions, we selected 4 uncorrelated conditions from Nguyen Ba *et al*. (16) as “hub” environments. We then chose 5 perturbations, which we add to the base media of each of the four hub environments, each at several different levels. We also chose an additional 5 perturbations which we applied at several different levels to a single hub environment. In total, this led to a set of 101 environments, summarized in Table 1 (see Table S1 for a complete list).

Our choice of specific hubs and perturbations was motivated by practical considerations and by previous work analyzing similar conditions. For example, several of the perturbations are similar to those analyzed in previous studies (4, 16, 24, 25), and collectively affect a broad range of cellular processes. We performed pilot experiments using a smaller set of F1 genotypes to choose the exact levels of each perturbation for each hub; our goal was to span a wide range of phenotypic correlations across each perturbation gradient, from highly correlated to the hub environment at the lowest perturbation level to substantially lower correlations with the hub environment at the highest perturbation level.

### Measuring growth rate phenotypes

To measure relative growth rate phenotypes (i.e. the competitive fitness of each segregant relative to the average) in a given media environment, we used bulk competition assays followed by barcode sequencing and fitness inference (Fig. 1B,C), following the protocol described in Nguyen Ba *et al*. (16) with minor modifications. Briefly, we pool all *∼*100,000 F1 segregants into a single flask and propagate this pool in batch culture for 8 days with daily 1 : 2^6^ dilutions (6 generations per day). At generations 6, 12, 30, 42, and 48 we conduct amplicon-based sequencing of the barcode region, and use barcode read counts to quantify the frequency of each segregant at each timepoint. We jointly infer the competitive fitness of all segregants in the pool simultaneously from these frequency trajectories using the method described in Appendix 2 of Nguyen Ba *et al*. (16), but with minor modifications. These bulk assays and fitness inference methods are described in more detail in Supporting Text 1 and Supporting Text 2 respectively.

Using this approach, we estimated the fitness of our *∼*100,000 segregants in each of our 101 different environments. We conducted all fitness assays in 2 independent replicates. In total, this effort involved more than 20 million fitness measurements across different genotypes, environments, and replicates. The accuracy of the resulting fitness measurements is primarily limited by depth of the barcode sequencing. In this study, average depth was *∼*80x (per barcode per timepoint per environment per replicate), resulting in a mean fitness correlation between replicates of *R*^2^ = 0.726 (Fig. S1).

### Phenotypic correlation structure reflects environmental structure

To analyze the degree of similarity in the genotypephenotype landscapes across our 101 environments, we first calculated the correlations in segregant fitness in each pair of environments (Fig. 1D). We find that the resulting correlation matrix has a complex structure that reflects several aspects of the relationships among environments. For example, we see 4 large blocks of relatively high correlation, each corresponding to a different hub environment (‘Gu’, ‘Li’, ‘YNB’, and ‘YPD’ in Fig. 1D). Within each of these large correlation blocks, we see smaller correlation blocks corresponding to environments with the same perturbation (with correlations typically stronger among environments with similar levels of that perturbation). Finally, there is also a more subtle trend of increased correlations between environments with the same perturbation level, but different hub environments (Fig. 1E, F). While there are also a variety of other subtle features of this correlation matrix, these results show that broadly speaking the hub-and-perturbation structure of the environments we analyze does leave the expected signatures in the structure of the fitness correlation matrix.

### Inferring pleiotropic genotype-phenotype maps

We next aimed to infer the genetic architecture of each of our 101 growth rate phenotypes by exploiting associations between the genotypes and phenotypes of the segregants in our mapping panel. Because our segregant panel is designed for this purpose, we can do so using standard methods from QTL mapping. However, this inference involves two key challenges relevant for our goals in this study.

First, standard methods in QTL mapping often first look for associations between genotype at each locus and a given phenotype using methods based on simple linear regression. For example, one can compute a log-odds (LOD) score describing the likelihood that a given locus is associated with a phenotype. However, because loci are typically tightly linked across substantial genetic distances (this is particularly true in our F1 cross), large regions of the genome surrounding a true causal locus will show significant LOD scores. In large segregant panels such as ours, nearly the entire genome is often statistically associated with a given phenotype, and identifying the causal loci and their effect sizes becomes difficult.

The second major challenge is that most existing methods for QTL mapping treat each phenotype independently. These approaches are poorly suited to our central goal of inferring the structure of pleiotropy across many correlated phenotypes. The central problem is that, because of genetic linkage, a single pleiotropic locus that affects multiple phenotypes may be mapped to separate, linked loci in each phenotype if these phenotypes are treated separately in our inference of the genotype-phenotype maps.

Recent work by Wang, Zou, and colleagues (26–28) introduced the SuSiE and mvSuSiE algorithms, which address the challenges described above. These algorithms use a Bayesian statistical framework to find sparse solutions to linear regression, naturally yielding both best-estimate values and uncertainties for QTL positions and effect sizes, and in the case of mvSuSiE also infer the genetic architecture of multiple phenotypes simultaneously. However, this approach is computationally expensive and proved intractable given the size of our dataset. We therefore instead implemented a multiphenotype extension of the method used by Nguyen Ba *et al*. (16), which extends their forward search procedure to jointly map the genetic basis of multiple phenotypes, as described below.

### Joint forward search algorithm

The genotype-phenotype map for each trait can in general be written in the form

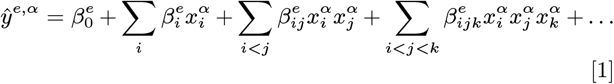

where *ŷ*^*e,α*^ is the predicted phenotype *e* (i.e. relative growth rate phenotype in that environment, so *e* equivalently indexes the environment) for segregant *α*, 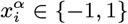 is the genotype of segregant *α* at locus *i*, 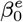 is the average value of phenotype *e* across all possible genotypes, 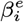 is the effect of locus *i* on phenotype *e* averaged over all genetic backgrounds at other loci, 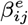 is the pairwise epistatic interaction between loci *I* and *j* on phenotype *e* averaged over all backgrounds at other loci, 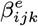 is a three-way epistatic interaction term, and so on. Our goal is then to use the segregant genotypes and measured phenotypes to infer the *β*^*e*^ terms.

Because the number of possible genotypes in any mapping study is always much larger than the sample size, we cannot fit the full genotype-phenotype map in Eq. (1). Here, we follow the typical practice of first fitting an additive model to the data (i.e. setting all epistatic terms to 0). Because even the number of potential additive terms (41,594; the number of SNPs that differentiate parental strains) approaches the size of our segregant panel, we use regularized linear regression to avoid overfitting. This identifies a relatively small number of causal loci that have nonzero effect on phenotypes, with all other 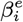 terms being set to 0. We then add potential pairwise epistatic terms, considering only interactions among loci inferred to have a nonzero additive effect (and with additional regularization to avoid overfitting, which sets most potential epistatic terms to 0). While third and higher-order interactions may also be present, we lack statistical power to infer these effects even given the large sample size of our segregant panel.

To implement this regularized inference procedure, we define a cost function:

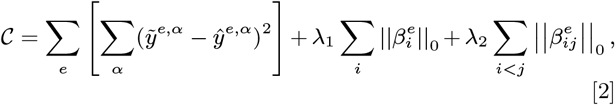

where 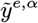 is the measured phenotype *e* for segregant *α* and *λ*_1_ and *λ*_2_ are regularization coefficients that penalize the number of nonzero additive 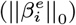 and epistatic terms 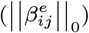 in the model respectively. Note that we explicitly allow for different regularization coefficients for additive and epistatic terms. However, we do not regularize our model across environments. In other words, once we identify a locus as causal, there is no additional penalty for it to have a nonzero effect size across more phenotypes. This choice not to regularize across environments is made in part for technical reasons: our optimization algorithm becomes computationally intractable when QTLs are not fully shared across environments. This ensures that all environments involve the same set of causal loci (though with independent effect sizes), facilitating cross-environment comparisons between models. However, it is important to recognize that this is a choice which favors the identification of pleiotropic loci. While the model can and does find that some loci are less pleiotropic than others (with very small effect sizes inferred in environments where that locus does not contribute substantially), it is in some sense at the opposite extreme from finding causal loci independently in each environment.

We note that, following Nguyen Ba *et al*. (16), we chose to use an *L*_0_ norm for regularization, rather than using Lasso (*L*_1_) or Ridge (*L*_2_) regression. This is because these latter methods perform poorly when predictors are highly collinear (29), as is the case in this study due to the relatively sparse recombination in our F1 panel. In particular, they tend to distribute predictor effect sizes between the correct position and spurious collinear linked positions. However, exact optimization of this cost function with the *L*_0_ norm would require fitting every possible additive model — in other words, every combination of nonzero QTLs — a process that is NP-hard (30) and intractable for the size of our dataset. Instead, we approximately optimize following the forward-search algorithm of Nguyen Ba *et al*. (16).

This forward-search algorithm is a greedy procedure that iteratively adds new QTLs to the model until a stopping criterion is met. In the first step, we regress every SNP in our panel against all 101 phenotypes, allowing each locus to have an independently inferred effect size in every environment. We find the locus that minimizes the summed squared residuals across all environments (i.e. the locus that minimizes Eq. (2)) and add it to the model, with the same position but independent effect sizes across environments. In the next step, we again search for the locus that minimizes Eq. (2) while allowing the effect sizes (but not position) of the previously added locus to be refit. We repeat the procedure until adding new QTLs no longer improves the regularized cost function. At times, the forward search procedure incorrectly assigns one large-effect QTL to a region that is better described by two (or more) small-effect QTLs. To allow the algorithm to re-optimize QTL locations when new QTLs are added in their vicinity (which effectively makes it possible to split largeeffect QTLs into multiple smaller-effect ones), we refit all QTL positions at the end of every fifth QTL addition step. Thus, one by one, we allow already added QTLs to find a new optimal position in a region bounded by neighboring QTLs or a chromosome boundary, whichever is closer. We note that because our ability to distinguish between the effects of tightly linked loci is limited, we often do not have single-nucleotide resolution on the identity of the true causal allele underlying each QTL; this is particularly true for smaller-effect loci, where statistical power is inherently more limited. For each QTL, we therefore identify both the “lead SNP” (i.e. the locus identified as optimal according to the algorithm described above), as well as a confidence interval around this lead SNP, defined as the region in which assigning the QTL to any SNP in that region (instead of the lead SNP) would increase our cost function by at most *λ*_1_. Intuitively, this can be thought of as the set of SNPs that we are as confident contains the true position of the QTL as we are that no new QTLs should be added to the model.

Despite using *L*_0_ regularization, the collinearity in our dataset led to some artifacts in the results of our forward search. Specifically, when nonzero terms are added at two tightly linked loci, their effect sizes can become unconstrained and they are assigned unrealistically large opposite sign values. Since we believe such values are far more likely to reflect artifacts than biological reality, we collapse these tightly linked large-effect, opposite-sign QTL pairs into a single QTL whose effect size we re-infer. In practice, this typically involves fewer than 5 pairs of QTLs and tends to improve prediction accuracy on held-out data.

The choice of regularization parameter (*λ*_1_ here while fitting only additive coefficients) determines the number of QTLs that are added to the model. However, we have no *a priori* reason to set *λ*_1_ to any particular value. Therefore, we employ a nested cross-validation procedure to learn the best choice of *λ*_1_ from the data. We first randomly partition the data into 10 equally sized batches and save one for final model validation. With the remaining 90% of the data, we run our forward-search algorithm nine times, each time leaving out a different batch from the training data and stopping each run of the algorithm once 480 QTLs have been added (empirically, this is the point at which we are confident the models have begun overfitting). We find the *ex post facto* optimal value of *λ*_1_ for each iteration of the algorithm by finding the *λ*_1_ that would have maximized the predictive power of the model in the held-out batch in that run. We then re-pool all nine training batches and run the algorithm one final time, setting *λ*_1_ equal to the mean of the optimal values discovered from the sub-batch runs.

After inferring causal loci in our additive model, we add epistatic terms in a similar manner: at each step of the forwardsearch algorithm here, we add the pair of loci to the model that minimizes the summed-squared residuals, and again reoptimize the effect sizes but not positions of all linear and already added epistatic terms. We determine the regularization coefficient for epistatic terms *λ*_2_ in the same manner as for additive coefficients. As mentioned above, we only consider interaction terms between loci included in the additive model to reduce the model search space and allow computational tractability.

## Results

The joint QTL mapping approach described above yielded 382 quantitative trait loci (QTLs), each with inferred additive effect sizes in each of the 101 environments. We show these inferred genetic architectures, described by the QTL positions and effect sizes, for the cycloheximide perturbation across all four hub environments in Fig. 2A (see Fig. S2 for all 101 environments). In addition, we identified 921 pairwise epistatic interactions among these 382 loci, that also have independently inferred effect sizes in each of the 101 environments. We illustrate these interactions for five different cycloheximide concentrations in the YNB hub in Fig. 2B. The complete set of 382 loci (including lead SNP and confidence intervals), 921 pairwise interactions, and inferred additive and epistatic effect sizes and confidence intervals on QTL position, are provided in Table S2.

**Fig. 2.**
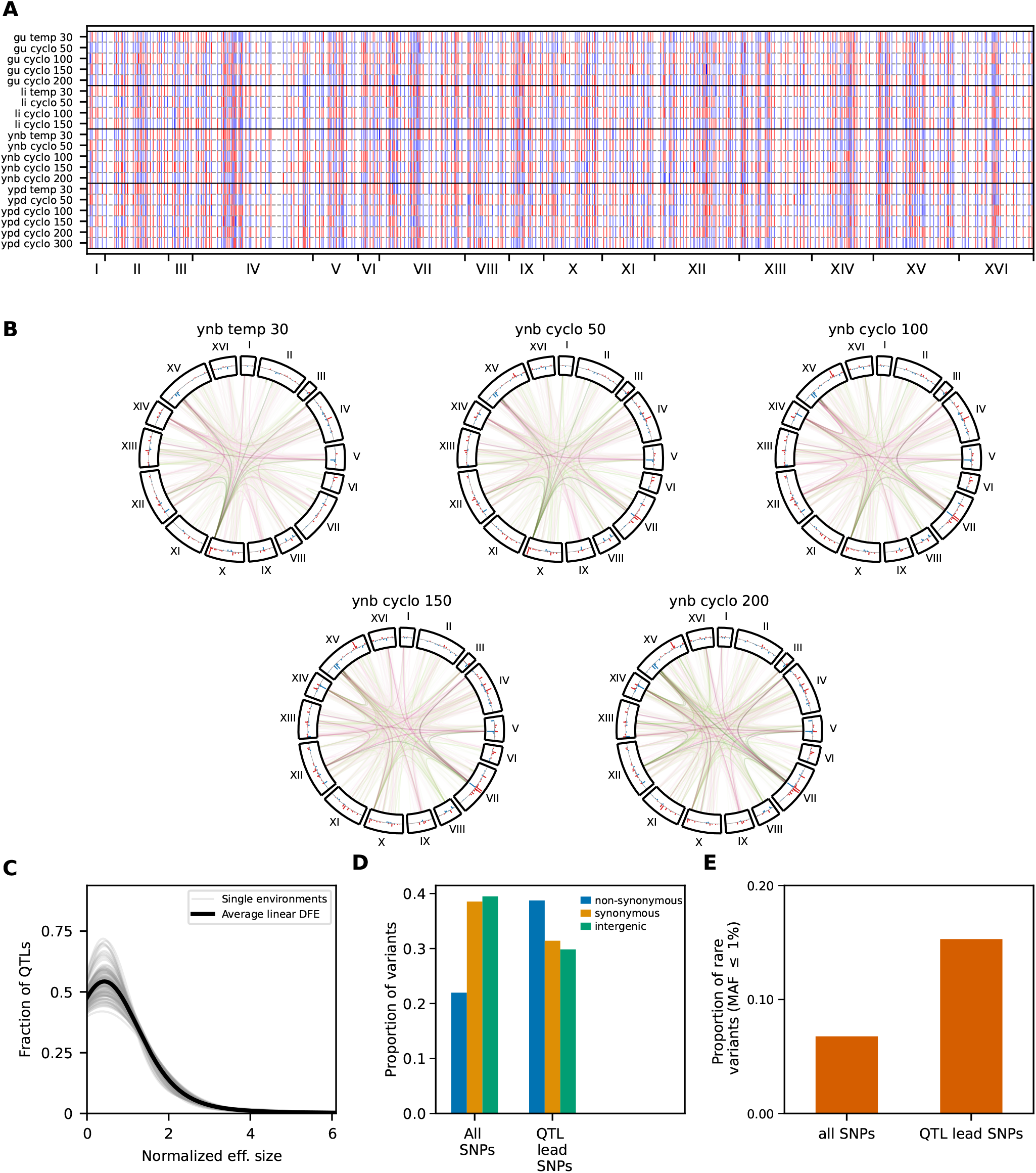
Summary of inferred genetic architecture. **(A)** QTL positions and inferred additive effect sizes in environments with a cycloheximide perturbation gradient. Environment names are shown along the y-axis. Loci are ordered according to genomic position (chromosome numbers indicated at bottom). The 382 inferred QTLs are highlighted with vertical bars, with inferred additive effect sizes shown by color intensity; red and blue colors correspond to cases where the RM or BY parental allele is more fit respectively. **(B)** Pairwise epistatic interaction network of fitness in the YNB hub, for five different cycloheximide perturbation concentrations (0, 50, 100, 150, and 200 nM). The 382 QTLs are arranged by genomic position around the circle with the additive effects being represented by the bars above and below the circle; curved lines connecting QTLs indicate inferred epistatic interactions. Color intensity reflects the epistatic effect size, with pink and green colors denoting negative and positive epistasis respectively. Negative epistasis (pink) is when the more fit allele at a locus has a smaller effect when the more fit allele is present at the other locus as well and vice versa for positive epistasis (green). **(C)** Distribution of additive effect sizes for individual environments (light gray, normalized by the mean effect size in that environment) and averaged across all environments (black). **(D)** The proportion of non-synonymous, synonymous, and intergenic variants among all SNPs and QTL lead SNPs. **(E)** The proportion of all SNPs and QTL lead SNPs that are found at low frequency (minor allele *≤* 1%) across the set of 1,011 natural yeast isolates from the yeast genomes collection (21).

### QTL effect sizes are exponentially distributed

We find that the distribution of the absolute value of the additive QTL effect sizes is roughly exponential in all 101 environments, except at very small effect sizes (Fig. 2C). The observed drop in the distribution at these small effects is likely because we are underpowered to detect small-effect QTLs below our limit of detection. This suggests that increasing the size of the segregant panel would lead to further increases in the number of small-effect loci identified. Similarly, we found that the distribution of epistatic effect sizes is also roughly exponential, except at very small effect sizes below our limit of detection (Fig. S3).

### QTL lead SNPs are enriched for non-synonymous and rare variants

We next sought to analyze general patterns in the identities of loci mapped using our joint forward search. Despite the uncertainty in the identity of the true causal SNP at each QTL, we nevertheless observe that QTL lead SNPs are enriched for non-synonymous variants in coding sequence (see Supporting Text 3 for SNP annotations; *p <* 10^*−*13^; *χ*^2^_*df* =2_ test; Fig. 2D). This enrichment is consistent with previous work (16), and may reflect the fact that non-synonymous variants are more likely to have measurable effects on our growth rate phenotypes than synonymous or intergenic variants. We also find that the mean QTL effect size across all 101 environments is larger for QTLs with non-synonymous lead SNPs than for QTLs with synonymous or intergenic lead SNPs (*p* = 0.0178 and *p* = 0.0549 respectively, two-sided Mann-Whitney U test; Fig. S4). This trend persists when using the maximum effect size of a QTL across all 101 environments in place of the mean effect size, though in this case the p-values are larger and the trends are not significant (*p* = 0.2822 and *p* = 0.4218 when comparing non-synonymous lead SNPs to synonymous and intergenic respectively). In contrast, the mean and maximum effect sizes of QTLs with synonymous lead SNPs are similar to those for QTLs with intergenic lead SNPs.

We find a significant enrichment of QTL lead SNPs among variants that are at low minor allele frequency (≤1%) in the 1,011 strains from the yeast genomes collection (21) (*p <* 10^*−*9^; *χ*^2^_*df* =1_ test; Fig. 2E). There is no overall relationship between QTL effect sizes and minor allele frequency (Fig. S5; *p >* 0.2 for the Spearman rank correlation). However, alleles with very low minor allele frequency have on average larger effect sizes (red points in Fig. S5). These results are consistent with recent work suggesting that rare, recently-arisen deleterious alleles contribute significantly to growth rate phenotype variation in budding yeast (14, 16, 31).

### Gene ontology enrichments among QTL lead SNPs

We next looked for functional relationships using gene ontology term (GO-term) enrichment (32–34) among the genes containing QTL lead SNPs (Supporting Text 4). We found that a total of 74 GO terms were significantly enriched in our dataset at an FDR threshold of 0.05 (Table S2). In Table 2, we show the 12 significantly enriched GO terms that are the most “derived” (i.e. they have no significant “child” GO terms). We see that these QTL lead SNP genes are highly enriched for transcription factors and other genes involved in transcriptional regulation, starvation response, TOR signaling, DNA replication, and cell cycle control, among several others.

**Table 2.**
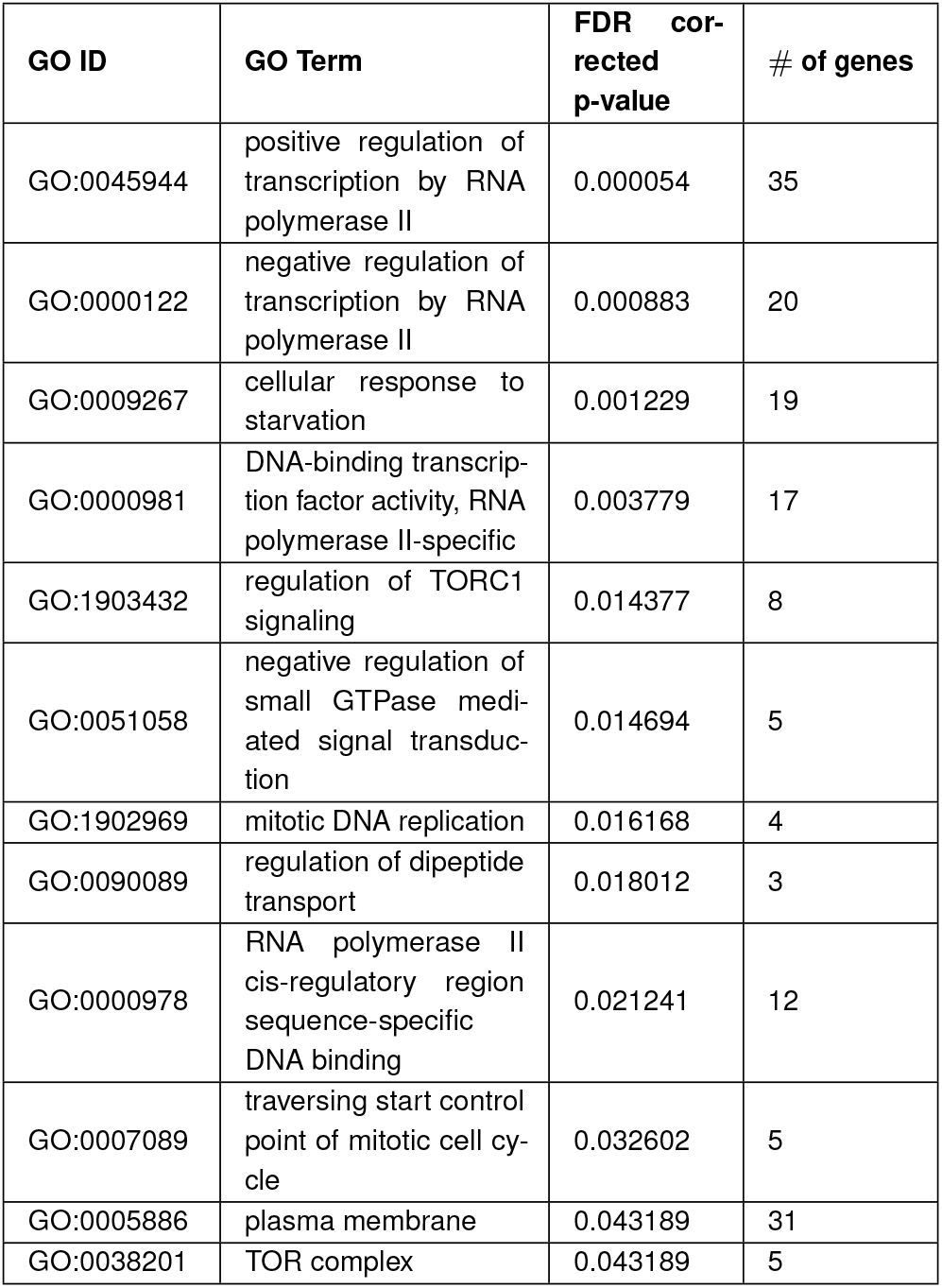
Significantly enriched derived GO terms.

| **GO ID** | **GO Term** | **FDR cor-rected p-value** | **# of genes** |
|---|---|---|---|
| GO:0045944 | positive regulation of transcription by RNA polymerase II | 0.000054 | 35 |
| GO:0000122 | negative regulation of transcription by RNA polymerase II | 0.000883 | 20 |
| GO:0009267 | cellular response to starvation | 0.001229 | 19 |
| GO:0000981 | DNA-binding transcrip-tion factor activity, RNA polymerase II-specific | 0.003779 | 17 |
| GO:1903432 | regulation of TORC1 signaling | 0.014377 | 8 |
| GO:0051058 | negative regulation of small GTPase medi-ated signal transduc-tion | 0.014694 | 5 |
| GO:1902969 | mitotic DNA replication | 0.016168 | 4 |
| GO:0090089 | regulation of dipeptide transport | 0.018012 | 3 |
| GO:0000978 | RNA polymerase II cis-regulatory region sequence-specific DNA binding | 0.021241 | 12 |
| GO:0007089 | traversing start control point of mitotic cell cy-cle | 0.032602 | 5 |
| GO:0005886 | plasma membrane | 0.043189 | 31 |
| GO:0038201 | TOR complex | 0.043189 | 5 |

### Gradient-ordered and “U-shaped” patterns of effect size are common

Next we sought to understand how the QTL effects across environments are jointly determined by the hubs and perturbations. We first analyzed how QTL effect sizes varied across perturbation gradients in environments with the same hub. If a locus is involved in the response to a given perturbation, we might expect the absolute effect size of the corresponding QTL to increase along the perturbation gradient, and for this same pattern to be seen across all hubs. We find that many QTLs do in fact show a “gradient-ordered” pattern: their effect size varies monotonically (strictly increases or decreases) as the perturbation gradient level increases. For example, Fig. 3A illustrates the variation in effect sizes of the thirty QTLs with strongest (average) absolute effects in the YNB hub environment across the cycloheximide perturbation; sixteen of these are gradient-ordered (left panel) while the remaining fourteen are not (right panel). Nearly every hubperturbation combination exhibits a significant excess of such gradient-ordered QTLs (*p <* 10^*−*7^ in 23 of the 25 combinations and *p <* 0.05 in 24 of the 25; Fig. 3B).

**Fig. 3.**
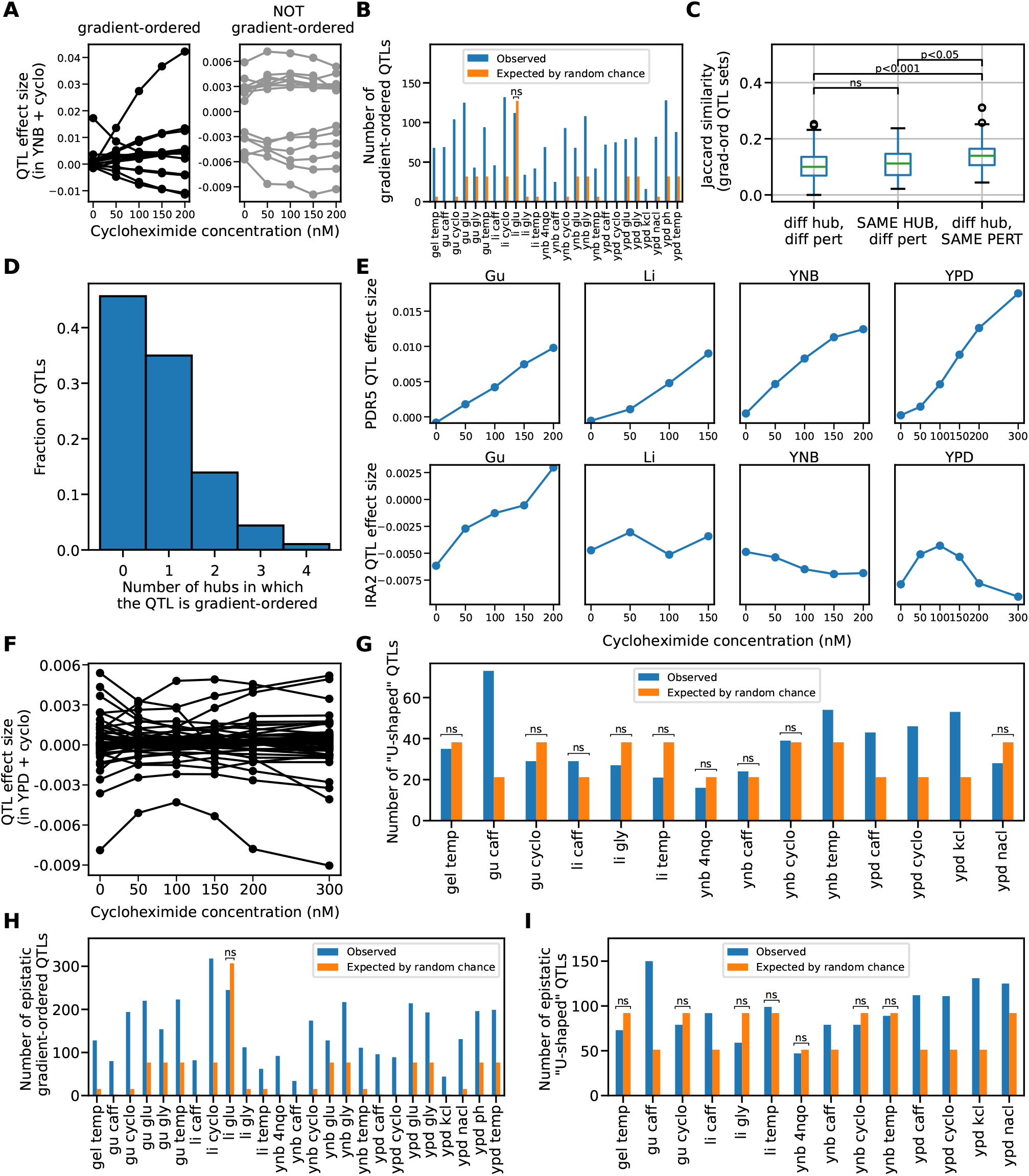
Gradient-ordered and U-shaped patterns in QTL effects. **(A)** Inferred effect sizes of the 30 strongest-effect QTLs in the YNB hub environment across the cycloheximide gradient. Gradient-ordered (16/30) shown at left, non-gradient-ordered shown at right (14/30). **(B)** Observed (blue bars) and expected (by permutation test; orange bars) number of gradient-ordered QTLs in each of the 25 hub/perturbation combinations. **(C)** Jaccard index of similarity between sets of gradient-ordered QTLs with different hubs and different perturbations (left), same hub and different perturbation (middle), and different hubs and the same perturbation (right). **(D)** Fraction of QTLs that are gradient-ordered across 0, 1, 2, 3 and all 4 hubs for the same perturbation (averaged over the five perturbations). **(E)** QTL effect size as a function of cycloheximide concentration in all four hub environments for two different QTLs, with lead SNP in PDR5 (top row) and IRA2 (bottom row). **(F)** Inferred effect size as a function of cycloheximide concentration in the YPD hub for the 46 QTLs with U-shaped patterns of effect size variation in this hub and perturbation. **(G)** Observed (blue bars) and expected (by permutation test; orange bars) number of U-shaped QTLs in each of the 14 eligible hub/perturbation combinations. **(H)** Observed (blue bars) and expected (by permutation test; orange bars) number of gradient-ordered inferred pairwise epistatic effects in each of the 25 different hub/perturbation combinations. **(I)** Observed (blue bars) and expected (by permutation test; orange bars) number of U-shaped pairwise epistatic effects in each of the 14 different eligible hub/perturbation combinations.

While a locus with gradient-ordered effects across a perturbation suggests that the locus is involved in a response to the perturbation, we observe that such gradient-orderings are not consistently found across all hubs. Only a small fraction of gradient-ordered QTLs were “hub-independent” (rightmost bar of Fig. 3D), meaning they showed gradient-ordered effect sizes across a given perturbation type in all four hub environments (as illustrated for the multi-drug resistance gene PDR5 in the top panel of Fig. 3E). Of the 20 QTLs that were found to be hub-independent, 14 of them are gradient-ordered in the cycloheximide perturbation, including several with lead SNPs in genes previously implicated in cycloheximide resistance (PDR1, YOR1, PDR5, TUP1, and SKI8 (36–38)). The vast majority of QTLs instead show more complex patterns of variation in effect size across our structured panel of environmental conditions (as illustrated for the IRA2 QTL across the cycloheximide gradient in the lower panel of Fig. 3E; see Table S3 for a comprehensive list).

To further understand these more complex patterns of variation, we quantified the similarity in sets of gradient-ordered QTLs between pairs of hub-perturbation combinations using the Jaccard Index (i.e. the number of shared elements of two sets divided by the number of elements in their union). We find that a gradient-ordered QTL is more likely to be gradientordered in a different hub with the same perturbation than be gradient-ordered in other types of perturbations (either with the same or different hub; *p* = 0.011 and *p* = 1.5*×*10^*−*4^ respectively, Mann-Whitney U test; Fig. 3C and Fig. S6A).

We next looked for QTLs that display a “U-shaped” pattern, with effect sizes that first increase, reach a maximum, and then decrease across a perturbation gradient (or vice versa). This pattern could for example reflect cases where one parental allele at a causal locus is optimal at some intermediate value of a perturbation. To reduce the number of such U-shaped QTLs we observe by chance, we require them to have at least 3 points supporting both the decreasing and the increasing phases. In Fig. 3F we show the U-shaped QTLs identified for the cycloheximide gradient in the YPD hub. We find a significant excess of these U-shaped QTLs in five of the fourteen eligible hub/perturbation combinations (those with at least five concentrations as required by our definition of U-shaped, Fig. 3G; *p <* 0.05, with *p <* 10^*−*4^ in four gradients, permutation test).

However, there is no clear pattern in sets of U-shaped QTLs shared across different hubs and perturbations as measured by the Jaccard Index and overlap coefficient (Fig. S6B,C). Thus there is no significant evidence of hub-independent patterns of effect size variation for U-shaped QTLs. However, we note that our power to detect these relationships is lower than for gradient-ordered QTLs due to the smaller number of gradients being considered and the lower number of U-shaped QTLs in each gradient.

### Epistatic and additive effects show similar patterns

As with additive effects, we observed a large excess of gradient-ordered epistatic effects in almost all perturbation gradients (Fig. 3H; *p <* 10^*−*7^ in 24/25 perturbation gradients). The number of gradient-ordered additive and epistatic effects are highly correlated across all our perturbation gradients (Fig. S7G; *R*^2^ = 0.715, *p <* 10^*−*6^). We also found a significant excess of Ushaped epistatic effect sizes in 7 of the 14 eligible perturbation gradients (Fig. 3I, *p <* 10^*−*3^ in all seven gradients), and the number of these U-shaped epistatic effects is correlated with the number of U-shaped additive effects (Fig. S7H, *R*^2^ = 0.503, *p <* 10^*−*2^). As with additive effects, very few epistatic effects are gradient-ordered or U-shaped in most or all hubs (Fig. S7C,F), further emphasizing that QTL effects are highly context (i.e. hub) dependent even when considering the same perturbation gradient.

As with additive effects, we found that gradient-ordered epistatic effects are more likely to be gradient-ordered in a different hub with the same perturbation than in a different hub with a different perturbation, though for epistatic effects this effect is only marginally significant (Fig. S7A,B, *p* = 0.054, Mann-Whitney U test). We found that U-shaped epistatic effects are similarly marginally more likely to be U-shaped in a different hub with the same perturbation (Fig. S7D,E; *p* = 0.055, Mann-Whitney U test); this effect was not significant in the additive case.

### Gradient-ordered QTLs are enriched for genes involved in corresponding biological processes

Our analysis of gradientordered QTLs was motivated by the hypothesis that they reflect a signature of underlying causal loci that participate in some biological process involved in response to the corresponding perturbation. To test this hypothesis, we asked whether QTLs with a lead SNP in genes (or in putative regulatory regions of genes) known to be involved in response to a given perturbation (as defined by phenotypic annotations from the Saccharomyces Genome Database, SGD (39)) are more likely to be gradient-ordered in that perturbation than QTLs with a lead SNP not located in such a gene. For the purposes of this analysis, we focused on the cycloheximide, 4-NQO, and caffeine gradients (these are the cases where genes could be most confidently assigned to be related to the perturbation). We find that the probability a QTL with lead SNP in a known cycloheximide or 4-NQO resistance gene is gradient-ordered (across at least 3 hubs for cycloheximide) in the corresponding perturbation is significantly higher than QTLs with lead SNPs in other genes (*p* = 0.08 and *p* = 0.044 respectively, one-sided Fisher’s exact test; Fig. 4A,C). Similarly, the change in inferred effect sizes across each perturbation gradient is larger for QTLs with lead SNPs located in the corresponding resistance genes (*p* = 0.009 and *p* = 0.008 for cycloheximide and 4-NQO respectively, one-sided Mann-Whitney U test; Fig. 4B,D). Similar results hold for the caffeine perturbation as well (Fig. S8).

**Fig. 4.**
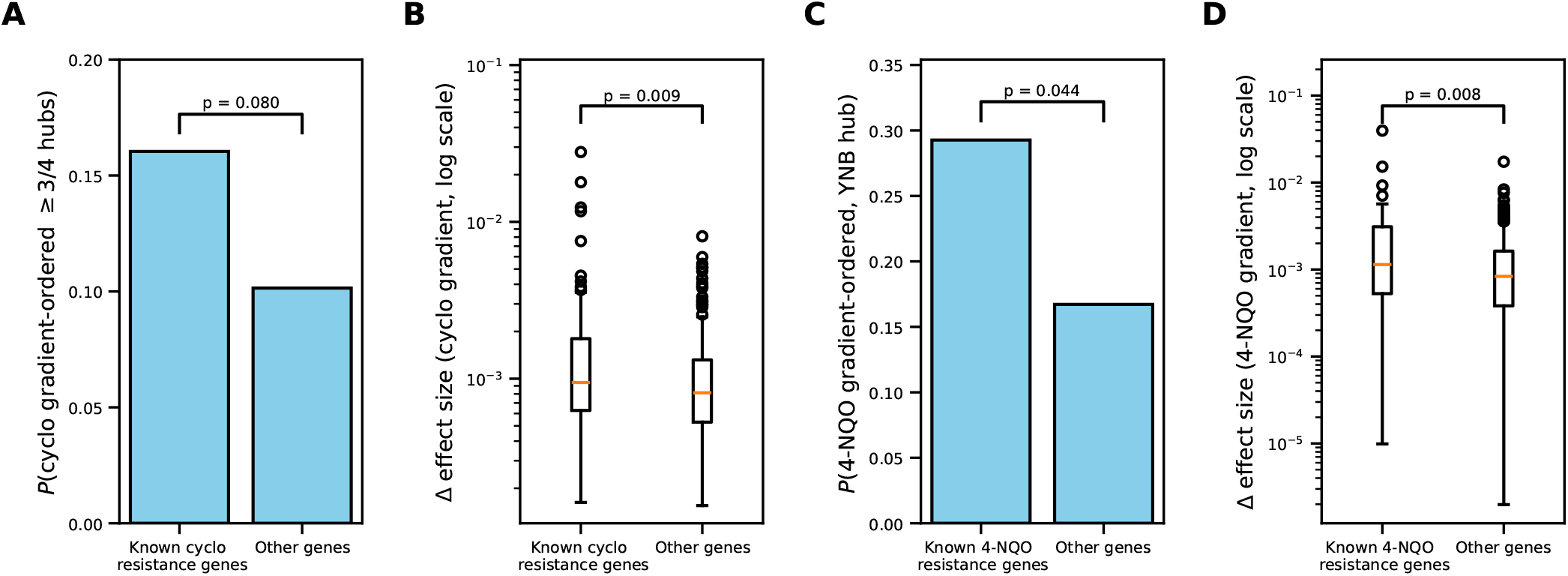
Enrichments for functionally related genes among gradient-ordered QTLs. **(A)** Fraction of QTLs in known cycloheximide resistance genes (versus in all other genes) that are gradient-ordered in the cycloheximide perturbation across at least 3 different hub environments. **(B)** Change in QTL effect size across the cycloheximide gradient for QTLs located in known cycloheximide resistance genes compared to for QTLs located in other genes. **(C)** Fraction of QTLs in known 4-NQO resistance genes (versus in all other genes) that are gradient-ordered in the 4-NQO perturbation. **(D)** Change in QTL effect size across the 4-NQO gradient for QTLs located in known 4-NQO resistance genes compared to for QTLs located in other genes.

### Pleiotropic patterns reflect the structure of genetic networks

To further investigate patterns of pleiotropy and probe their potential relationship with gene function, we computed the squared Pearson correlation (*R*^2^) of z-scored effect sizes across all 101 environments for all 382*×*382 pairs of inferred QTLs. We then hierarchically clustered the resulting correlation matrix (Fig. 5A). We immediately observe lower-dimensional structure in this matrix, with numerous blocks of highly correlated QTLs. To quantify this structure, we grouped the 382 QTLs into 118 different clusters based on this *R*^2^ metric of similarity between their normalized effect sizes (see Supporting Text 5 for details).

**Fig. 5.**
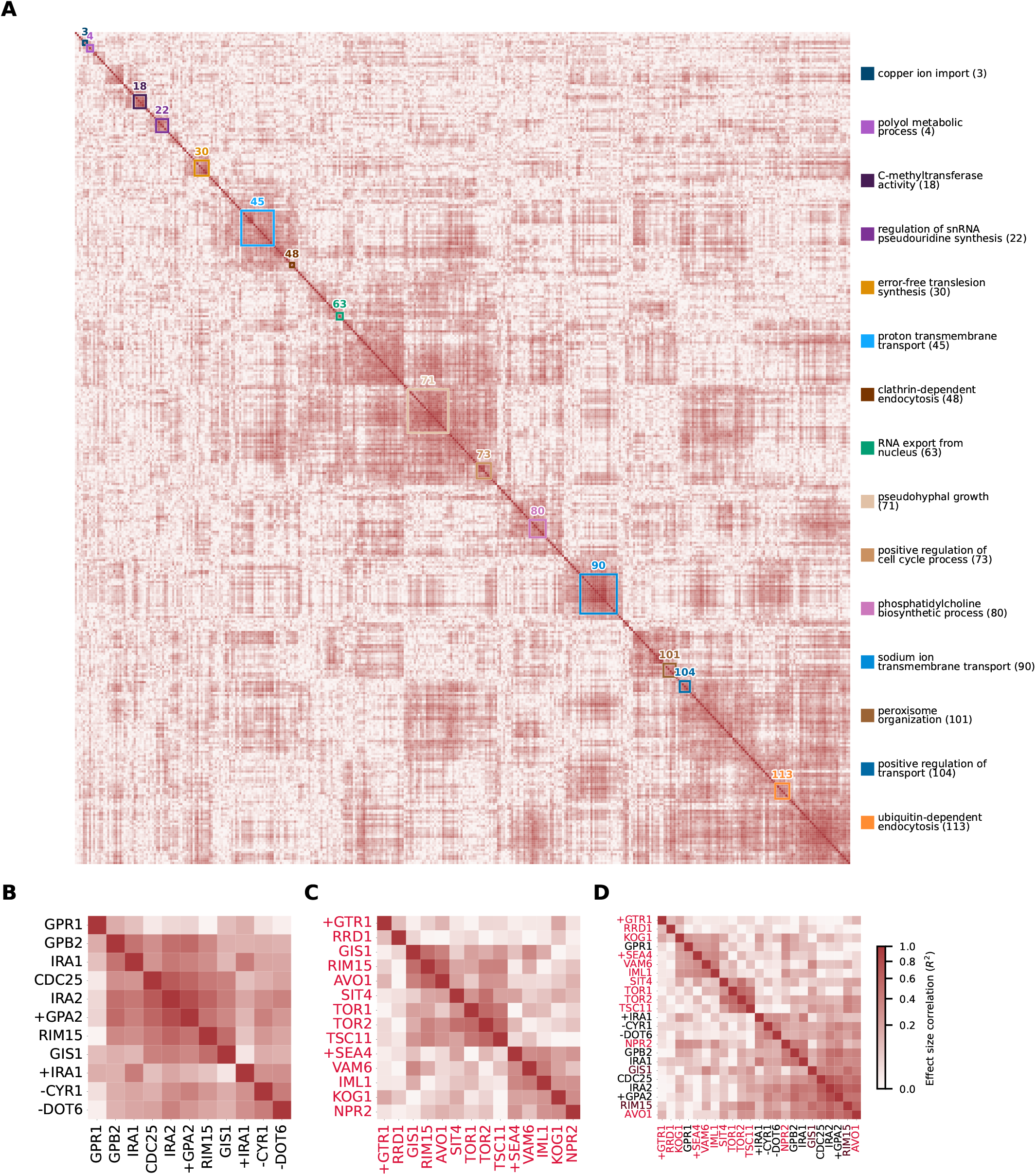
Correlations in effect sizes of inferred QTLs across environments. **(A)** Correlation across environments in (normalized) inferred additive effect size for all 382 *×* 382 pairs of QTLs, arranged by hierarchical clustering (ordered list of QTLs and corresponding lead SNP genes and/or lead intergenic regions shown in Table S5). QTLs are clustered into 118 different groups for downstream analysis. 16 of these clusters show a significant enrichment of QTLs associated with the same gene ontology term and are highlighted with colors shown on the right (with the exception of one cluster that does not have a significant GO term with fewer than 25 associated QTLs). **(B)** Heatmap showing the correlation in (normalized) fitness effect size for QTLs in the Ras/PKA pathway, **(C)** TOR pathway, and **(D)** both Ras/PKA (black labels) and TOR (red labels) pathway QTLs, with QTLs in genes shared across both pathways highlighted with brown labels.

To test whether this clustering based on pleiotropic patterns reflects the structure of genetic networks, we analyzed whether any gene ontology (GO) term was enriched among the genes containing lead SNPs for the QTLs in each cluster (compared to the full “background” set of 382 QTLs). For each cluster, we computed *p*-values for this enrichment (righttailed hypergeometric test) and performed multiple-testing correction using the Benjamini-Hochberg method (with false discovery rate FDR=0.05). We only considered GO term enrichment hypotheses that were driven by more than one QTL in a given cluster. The resulting corrected *p*-values for all hypotheses are shown in Table S5; see Supporting Text 5 for details.

Based on this analysis, we find 16 clusters that show a significant enrichment for the presence of 89 total GO terms. For each cluster, we highlight one of these significantly enriched GO terms in the legend to the right of Fig. 5A (with one exception; see caption); these representative GO terms are the most “derived” Biological Process GO terms corresponding to each cluster that have fewer than 25 QTLs associated with them in our dataset (25 is the number of QTLs in the largest of the 118 clusters and is thus a natural upper bound here). Cluster 18 has no significant Biological Process GO term and thus we show its only significant GO term instead (which is a Molecular Function GO term). We highlight here four of these clusters that contain large-effect QTLs where our inferred confidence interval around the lead SNP is small (*≈*3-4 genes). First, cluster 22 is annotated as “regulation of snRNA pseudouridine synthesis” but the more relevant function term here is TOR signaling (note that TOR signaling regulates starvation-induced pseudouridine synthesis (40)). Cluster 22 contains three genes involved in TOR signaling (TOR1, TOR2, and TSC11) along with AIT1/YDL180W, which is not GO annotated as a TOR signaling gene but has a known negative regulatory effect on TOR (41). Second, cluster 30 contains 3 genes (RAD5, REV1, and REV7) that are involved in error-free translesion synthesis (i.e. DNA damage repair). Finally, we identify two clusters with genes involved in salt/cation tolerance: cluster 90 contains ENA1, ENA5, and PHO89, three genes that are involved in transporting cations across the cell membrane, and cluster 104 contains SAT4, SKY1, and HAL5, three genes that are involved in maintaining cellular homeostasis for cations (annotated more generically as “positive regulation of transport”). Note that HAL5 contains two different lead SNPs corresponding to two different QTLs in our dataset; one QTL is found in cluster 90 (with ENA1 and ENA5) and the other QTL is found in cluster 104 (with SAT4 and SKY1). In summary, we are able to capture known biological relationships among multiple clusters and QTLs using this approach, which does not make use of any functional information about the genes involved.

These results show that clustering based on correlations in effect sizes across environments can identify signatures of the structure of genetic networks in yeast. In addition to this “top-down” analysis, we also explored a “bottom-up” approach, where we looked at effect size correlations between QTLs with lead SNPs in genes within the same biological pathway. As an example, we analyzed effect size correlations between QTLs in the Ras/PKA and TOR pathways (Fig. 5B-D). Both pathways are significantly overrepresented among our QTLs (Table 2); they are broadly involved in nutrient sensing and response, with the Ras/PKA pathway sensing and responding to glucose (i.e. carbon source) levels and the TOR pathway sensing and responding to amino acid (i.e. nitrogen source) levels. They are also a primary source of large-effect adaptive mutations in yeast experimental evolution experiments (42–44). Variation in genes within these pathways is common among natural yeast isolates, and much of the variation in these pathways appears to have functional effects and shows signs of local adaptation to specific environmental niches (45).

We find that the squared Pearson correlation (*R*^2^) of normalized effect sizes across our environmental conditions is much higher than expected by random chance between pairs of QTLs in the Ras/PKA pathway (Fig. 5B; *p <* 10^*−*4^, permutation test), pairs of QTLs in the TOR pathway (Fig. 5C; *p* = 1.34*×* 10^*−*2^), and the combined set of QTLs across both pathways (Fig. 5D; *p* = 5 *×*10^*−*4^). In other words, as expected, QTLs within the Ras/PKA and TOR pathways have increased effect size correlations compared to random QTLs.

We next investigated the finer structure in these effect size correlations. We find that Ras/PKA QTLs largely form a single cluster of high effect size correlation (with the exception of the GPR1 QTL; see Fig. 5B). This is in contrast with the TOR pathway shown in Fig. 5C, which contains at least 2 different groups: group 1 containing TORC1/TORC2 complex genes (TOR1, TOR2, TSC11, AVO1) and associated downstream response genes (GIS1, RIM15, SIT4), and group 2 containing genes upstream of TORC1/TORC2 that reside on the vacuolar and lysosomal membranes and detect amino acid levels (SEA4, VAM6, IML1 and NPR2) (46). Thus broadly speaking this clustering within the TOR pathway recapitulates its organization into “upstream” amino acid sensing components and “downstream” TORC1/TORC2 complexes and response regulators (though we note that the QTL in KOG1, a gene in the TORC1 complex, does break this pattern). We also note that the GTR1 and RRD1 QTLs form their own group among the TOR pathway genes; the simplest explanation for these “outliers” (GTR1 and RRD1 QTLs in the TOR case and GPR1 in the Ras/PKA case) is that the true causal variant is located in another gene within the confidence interval of the QTL and thus are included in this pathway analysis as false positives due to mapping errors.

As described above, the Ras/PKA and TOR pathways are largely thought to sense and respond to glucose and amino acid levels respectively. However, there is also documented “cross-talk” between these pathways (46, 47): they have been known to respond to other types of nutrient constraints and stresses and share some downstream signaling proteins (e.g. Rim15 (47)). To probe for signatures of this cross-talk in our data, we plotted the correlations of effect sizes for pairs involving both Ras/PKA and TOR pathway QTLs (Fig. 5D). We see several clusters in this data, which largely correspond to pathway identity, with some exceptions. However, we also see cross-cluster correlations between Ras/PKA and TOR pathway QTLs, consistent with the existence of cross-talk between these pathways that leaves visible signatures in the patterns of pleiotropy in our cross.

## Discussion

In this study, we measured the fitness of 100,000 F1 segregants in 101 different media environments arranged in a “hub-andperturbation” structure constructed by varying the strength of a variety of perturbations across a gradient in each of several different “hub” growth media. This produced an extensive map of the correlation structure of fitness across environments. We then developed a new joint QTL mapping method to use this data as the basis to map the fitness effects of 382 QTLs in each of these conditions.

We found that the genetic architecture of fitness in our dataset was very highly pleiotropic, with just 382 fitness QTLs being mapped across 101 different environments (compared to *∼*1800 QTLs that were independently mapped across just 18 environments in Nguyen Ba *et al*. (16) using the same collection of F1 segregants). This reflects the choices involved in the mapping pipeline we used in this study, which favors the identification of pleiotropic loci and represents an opposite extreme to the approach of mapping QTLs independently in each environment.

Given the hub-and-perturbation environmental structure, we might expect that QTLs involved in response to a particular perturbation would display monotonically increasing (or decreasing) effect sizes across increasing strengths of that perturbation. However, while we did observe a large excess of such gradient-ordered QTLs in nearly all of our perturbation gradients, very few of these were gradient-ordered across that perturbation in multiple hub environments. In addition, we also found examples of gradient-ordered QTLs shared in a variety of patterns across different hubs and perturbations (e.g. a QTL that is gradient-ordered in caffeine in one hub and in cycloheximide in the same or different hub). Thus, overall, most QTLs display more complex patterns of pleiotropy involving interactions between hubs and perturbations.

We found that the QTLs whose patterns of pleiotropy suggested they may be involved in response to a given stress (e.g. those that are gradient-ordered in the corresponding perturbation) were enriched for genes involved in the corresponding biological pathways. This suggests that the observed pleiotropic structure contains information about the structure of genetic networks in yeast. Motivated by this, we clustered QTLs based on their patterns of pleiotropy and investigated the relationship between these clusters and potential biological functions. For example, we found clusters that roughly correspond to processes such as salt/cation tolerance, the Ras/PKA pathway, TOR signaling, and others.

In principle, one could use this signal to identify novel associations between genes and functional pathways, and to generate hypotheses about potential interactions and connections between biological processes. For example, we found that the fitness effects of QTLs were correlated across the Ras/PKA and TOR signaling pathways, suggesting “cross-talk” between them. In addition, we can generate hypotheses about how specific variant alleles may affect pathway function. In principle, one could further investigate these hypotheses using other data sets, such as existing single-cell RNA sequencing data from these F1 segregants ((48); (49) also uses a BY/RM cross). Future work along these lines has the potential to map effects of these naturally occurring genetic variants at different levels of biological organization (protein, protein activity and phosphorylation, metabolite, flux, and pathway output) to generate a more comprehensive molecular picture of the genotype-to-phenotype map.

Several recent studies (17, 18, 50) have used earlier data sets describing the pleiotropic effects of variants across multiple environments as the basis for computational methods to identify lower-dimensional latent structure in the genotypephenotype map. The implication of this work is that this lower-dimensional structure potentially reflects (and could possibly be used to infer) the structure of genetic networks. Our study here provides a large new data set that could be used as the basis for these and other types of structure discovery methods. This is an interesting topic for future work.

We note that while we see many QTLs in nutrient sensing and response pathways, we see relatively few QTLs in metabolic pathways such as glycolysis, TCA cycle, and adenine biosynthesis. This observation suggests a possible mechanism of molecular adaptation in yeast: they prefer to fix mutations in upstream regulatory pathways that can widely affect metabolism (such as Ras/PKA and TOR) rather than directly tinkering with metabolism by fixing mutations in individual metabolic pathways. While this proposed mechanism is in line with the results of some laboratory evolution (43, 44) and multi-omics experiments (51), more targeted experiments are required to directly test this hypothesis.

## Acknowledgements

We thank Alex Nguyen Ba, Katherine Lawrence, Caroline Holmes, Misha Gupta, and other members of the Desai lab for many useful conversations. We also thank the Bauer core facility at Harvard for assistance with sequencing. SG acknowledges support from the NSF-Simons Center for Mathematical and Statistical Analysis of Biology at Harvard (award number #1764269) for a fellowship award. This work was supported by grants R01-GM104239 and R35-GM161612 from the NIH. The computations in this paper were run on the FASRC Cannon cluster supported by the FAS Division of Science Research Computing Group at Harvard University.

