## Supplementary Information for "Joint QTL mapping reveals patterns of pleiotropy across structured environments in budding yeast"

#### This PDF file includes:

Supporting text
Figs. S1 to S8
Legends for Tables S1 to S5
Legends for Datasets S1 to S3
SI References

#### Other supporting materials for this manuscript include the following:

Tables S1 to S5
Datasets S1 to S3

### Supporting Information Text

### 1. Bulk fitness assays

We measured the fitness of the 100,000 F1 segregants in our mapping panel using bulk fitness assays similar to those described by Nguyen Ba et al. (1), with several modifications (Fig. 1). Briefly, we thawed a glycerol stock of all pooled segregants and grew in rich laboratory media (YPD) for 5 generations. We then inoculated 1.56 mL of saturated culture into a 500 mL flask containing 100 mL of the growth medium of interest. We measured the fitness in each environment using two independently propagated sets of flasks, providing a technical replicate of our fitness measurements. The list of all environments for which we carried out duplicate fitness assays can be found in Table S1; all environments (media) had a final glucose concentration of 2% w/v and a final ampicillin concentration of $100\mu g/mL$ unless otherwise noted, and all incubation was done at 30°C except as noted for the temperature perturbation. The pooled F1 segregants were allowed to grow for 24 hours while being shaken vigorously (170 rpm, 5.08 cm orbit diameter). This grown-up culture of F1 segregants was then diluted into fresh medium (1.56 mL of culture into 100 mL of fresh medium) and propagated as described here for an additional 7 days (6 generations per day). Aliquots of the culture were pelleted and frozen at −80°C at generations 6, 12, 30, 42 and 48. We conducted these assays in relatively large volumes of growth medium (100 mL) to avoid severe bottlenecks at each transfer (∼1000 cells per segregant transferred every day).

We then lysed and extracted DNA from each cell pellet, as described by Nguyen Ba et al. (1). We then used 10 µL of the extracted DNA ($\sim 100 - 500$ ng) as template to PCR the genomic barcode and amplify it for 3 cycles. The first PCR was used to add a set of multiplexing indices and unique molecular identifiers (UMIs) that allow for the detection of "jackpotting" events that can occur during PCRs. This is also why the first PCR only had 3 cycles of amplification: the small number of amplification cycles prevents the frequencies of each barcode from changing too drastically after UMI addition. Primers were then removed from the PCR product using left-sided size selection with SPRIselect beads. This cleaned-up PCR product was then put through a second PCR to add sequencing adapters and a second set of multiplexing indices; this PCR had the "normal" 35 cycles of amplification. This PCR product was then subjected to a left-sided size selection to remove the primers. The cleaned-up PCR products from each environment × timepoint × replicate of a fitness assay (1110 total PCR products) were pooled together with an equal weight of DNA from each PCR product. We note that these methods for sequencing library preparation are very similar to those described in Nguyen Ba et al. (1).

The resulting pooled library was sequenced on an Illumina Novaseq instrument using an S4 flowcell (with a small number of reads/samples being sequenced using a Novaseq SP flowcell) at an average coverage of 80x per barcode per environment per timepoint per replicate. This sequencing coverage (80x per barcode) is around half the sequencing coverage used in previous work ((1); 185x per barcode), which leads to reduced precision in the resulting fitness measurements. However, simulations from Nguyen Ba et al. (1) showed that the accuracy of inferred fitness and QTL effect size increased relatively slowly with increasing sequencing coverage, and even an average sequencing coverage of 80x was expected to yield inferred fitnesses that were sufficiently accurate for our purposes ($R^2$ between inferred and true fitness $\sim 0.7 - 0.9$ at 80x sequencing coverage per segregant). This motivated our choice of a target coverage of 80x per segregant for our extremely broad survey of the phenotype space.

Raw sequencing data was first cleaned: only the lines in each fastq file containing the sequencing read itself were retained and reads containing long stretches of 10 or more guanines were discarded as they likely correspond to sequencing errors caused by low-intensity signal. Reads from different lanes of the same flowcell were combined before proceeding further. The sequencing reads were then parsed: raw sequencing reads that match the template of a genomic barcode were identified and the multiplexing index and barcode information were extracted from such matching sequencing reads. The barcodes were next error corrected: the sequenced barcodes (from the fitness assay) were mapped to the list of ∼100,000 known barcode sequences in our F1 segregant pool. This error correction allows for 2 mismatches between the sequenced and known barcodes, to account for sequencing errors. These error-corrected barcodes were then index corrected: only barcodes with the correct combination of multiplexing indices were allowed to pass through this filtering step, removing erroneous sequencing reads that result from index hopping and other processes. The barcodes were then counted (the number of occurrences of each unique barcode is calculated) and these barcode counts were joined across the different timepoints of the same replicate of a particular environment. These barcode counts across time form the basis for our phenotype inference described in the next section.

### 2. Phenotype Inference

Assuming deterministic dynamics and that segregants only interact via competitive exclusion (i.e. there is no frequency dependence), we expect the frequency $f$ of segregant $i$ in replicate $r$ to evolve according to the coupled differential equations

$$\frac{\partial f_{i,r}}{\partial t} = f_{i,r}(y_i - \overline{y}_r), \qquad [1]$$

where $y_i$ is the fitness of lineage $i$ and $\overline{y}_r$ is the (time-dependent) average fitness of the population in replicate $r$: $\overline{y}_r = \sum_i y_i f_{i,r}(t)$. The solution to these equations is

$$f_{i,r}(t) = \frac{f^0_{i,r} e^{y_i t}}{\sum_j f^0_{j,r} e^{y_j t}}, \qquad [2]$$

where $f^0_{i,r}$ is the population fraction of lineage $i$ in replicate $r$ at the start of the fitness assay (time $t = 0$) and time is measured in generations.

Inferring the fitness of a lineage in an environment then reduces to the problem of finding the set of $y_i$ (fitness of lineage $i$) and $f^0_{i,r}$ (initial frequency of lineage $i$ in replicate $r$) that best explains the barcode frequency data. We optimize under a Bayesian statistical framework, applying a Gaussian prior to population fitness (this prior reduces artifacts that otherwise arise for very low fitness segregants where read counts are typically low and noise is correspondingly high). We assume that the sequencing read counts $N_{i,r}(t)$ for a lineage $i$ in a given replicate $r$ and time $t$ are multinomially distributed (note this does neglect overdispersion due to PCR jackpotting and other effects; see Nguyen Ba et al. (1) for more details). This defines the following statistical model:

$$
\begin{aligned}
f_{i,r}(t) &= \frac{f^0_{i,r} e^{y_i t}}{\sum_j f^0_{j,r} e^{y_j t}}, \\
y_i &\sim \mathcal{N}(\mu, \sigma^2), \\
\vec{N}_{i,r}(t) &\sim \text{Multinomial}\left(\vec{f}_{i,r}(t), N_{\text{tot},r}(t)\right).
\end{aligned}
$$

Here the initial frequency of each lineage $f^0_{i,r}$ is a free parameter (constrained only to sum to one across lineages). The mean and variance of the Gaussian prior are not known *a priori*; we iteratively tune these parameters, with each successive iteration setting the prior based on the mean and variance of the fitness distribution from the previous iteration. We also note that this fitness inference is invariant to a constant fitness translation (i.e. only fitness differences between strains are relevant). We must therefore choose an arbitrary 0-point for fitness. Here we choose to define the fitness of the segregant with the most total read counts in an environment as having fitness equal to zero in that environment, as this tends to reduce the overall noise of the fitness inferred across all segregants. As in Nguyen Ba et al. (1), we fit the parameters of this model (the initial frequency $f^0_{i,r}$ and the fitness $y_i$ for all ≈100,000 lineages) using a preconditioned nonlinear conjugate gradient algorithm (also known as "Hager-Zhang" (2)).

The process outlined above jointly infers a single fitness value for each lineage using data from both replicate fitness assays (with initial frequency of each lineage allowed to vary across replicates). To estimate the precision of these measurements, we also repeated this inference procedure using data from each replicate separately (Fig. S1). We find that the $R^2$ between replicate fitness measurements averaged 0.726 across our 101 environments (range of 0.467–0.829).

We initially performed bulk fitness assays in a set of 111 environments. However, 10 environments were excluded from the final dataset because they failed checks at various stages of the quality control process, leaving us with our final set of 101 environments used in all further analyses. The excluded environments are listed in Table S1. Two of these 10 excluded environments were so stressful that cell density crashed during our bulk fitness assays, making it impossible to estimate barcode frequencies through time. For these environments we have no fitness measurements at all. In the other eight excluded environments, we were able to measure fitness using our bulk assays. For five of these, we find that the correlation in fitness between replicates is very poor ($R^2$ less than 0.4; Fig. S1), primarily due to low read counts which likely resulted from problems in library preparation or other aspects of our sequencing pipeline. For the other three, we find that sequencing depth or segregant diversity was very low in one of the replicates, preventing accurate quantification of errors.

### 3. SNP functional annotations

We annotated all variants identified as QTL lead SNPs according to mutation type (synonymous, nonsynonymous, or intergenic), minor allele frequency as computed from the 1011 Yeast Genomes collection (3), gene identity and essentiality (for coding mutations), and potential regulatory effects (for intergenic mutations). We also annotated gene identities and potential regulatory effects for all SNPs within the confidence interval of each QTL. We provide a complete list of these annotations in Table S2.

To generate gene name annotations for coding mutations, we used the gff file created by Nguyen Ba et al. (1). Specifically, we annotated a SNP as in a gene if that SNP is located in a non-dubious gene feature of the gff (but not in an intronic region). We also recorded whether each annotated gene is essential and non-evolvable essential according to the definition of Liu et al. (4). We note that a SNP can be located in and thus annotated with multiple genes (this is rare, as gene overlap is rare in the *S. cerevisiae* genome). We classified coding SNPs as either synonymous or nonsynonymous, and all other SNPs as intergenic. For non-synonymous SNPs, we further classify them as nonsense SNPs if they create a premature stop codon.

We classified intergenic SNPs as potentially "upstream regulatory" for a gene if it was located within 500 bp upstream of the start of the gene's CDS and "downstream regulatory" for a gene if it was located within 500 bp downstream of the end ("stop" in the GFF file) of the gene's CDS. A given SNP can be (and often is) annotated as regulatory for multiple genes.

### 4. GO term annotation and GO enrichment analysis

We used the Python goatools library (5) to perform GO enrichment analysis. Our query gene set for this analysis is the 265 distinct genes corresponding to lead SNPs of one of our 382 identified QTLs. We compare this to a background set consisting of the 4183 genes that contained at least one SNP in our mapping panel. We identified enrichments for GO terms using "tier 1" GO annotations (those assigned based on evidence from low-throughput validation experiments; these are generally considered the most trustworthy annotations, and correspond to evidence codes IDA, IMP, IGI, IPI, IEP, EXP, TAS, and IC). GO annotations were propagated using the "True Path Rule" (meaning that a gene associated with a GO term is also associated with all ancestral GO terms; true path propagation was done using both "*is_a*" and "*part_of*" relationships).

We find that our query set of 265 genes is associated with 2303 unique GO terms (Biological Process [BP] + Molecular Function [MF] + Cellular Component [CC] terms) and our background set of 4183 genes is associated with 6704 unique GO terms (again, BP + MF + CC terms). We computed a $p$-value for the enrichment (and depletion) of each of these 6704 total GO terms in our dataset using GOEnrichmentStudyNS (which uses Fisher's exact test). We corrected these $p$-values for multiple testing using both a Benjamini-Hochberg FDR correction and a Bonferroni correction (applied separately to BP, MF, and CC GO terms). We find a total of 74 GO terms that are significantly enriched at FDR threshold of 0.05 and a total of 39 GO terms that are significant at a Bonferroni corrected $p$-value threshold of 0.05. A full list of these significantly enriched GO terms can be found in Table S2. We found no evidence for the depletion of any GO terms in our query gene set at a significance threshold of 0.05 (both FDR and Bonferroni).

For our analysis of the Ras/PKA and TOR pathways, we annotated QTLs with lead SNPs in these genes with additional GO terms corresponding to the different subcomponents of these pathways. Specifically, for the Ras/PKA pathway we considered three different subcomponents: negative regulators of Ras, positive regulators of Ras, and Ras/PKA pathway downstream effectors (transcription factors, kinases, and phosphatases). For the TOR pathway we considered eight subcomponents: the TOR complex 1 (TORC1), TOR complex 2 (TORC2), the EGO complex (EGOC), the vacuolar complexes SEACIT (SEA subcomplex inhibiting TORC1) and SEACAT (SEA subcomplex activating TORC1), other vacuolar genes affecting TOR signaling, the Sch9 branch of the TORC1 pathway, and the PP2A branch of the TORC1 pathway. The list of genes in these subcomponents was gathered from Conrad et al. (6), Venkataram et al. (7), Plank (8), and SGD. A full list of all of these Ras/PKA and TOR pathway subcomponents and their corresponding genes can be found in Table S5.

### 5. Hierarchical clustering of QTLs

We divided our 382 QTLs into clusters based on their inferred effects on our 101 phenotypes. To do so, we first built a distance matrix between all QTL pairs, where the distance between each pair is defined as $1 - r^2$, where $r$ is the Pearson correlation coefficient of their normalized (z-scored in each environment) effect size vectors. We then used this distance matrix to cluster QTLs using the UPGMA (Unweighted Pair Group Method with Arithmetic mean) algorithm (9). This algorithm starts with each QTL as a separate cluster, and then merges the two nearest clusters at each step until a single cluster remains, creating a dendrogram representing the hierarchical cluster structure. We can then divide the QTLs into a defined set of clusters by cutting the tree at a threshold depth in this dendrogram (we choose 0.7, so that roughly speaking QTLs are grouped into the same cluster if their average $r^2$ is at least 0.3). This resulted in a set of 118 clusters.

We next turned to estimating GO enrichments within these 118 clusters. To do so, we use a more liberal set of GO annotations than described in Supporting Text 4 above. Specifically, we now also consider liberal or "tier 3" GO annotations based on high-throughput experiments, phylogeny, or other evidence, and we also consider both coding SNPs as well as intergenic SNPs with putative regulatory effects. Note that both coding and intergenic SNPs (with putative regulatory effects) were also considered for the analysis in Fig. 4. For each cluster, we tested for GO term enrichment. We note that to reduce multiple hypothesis testing corrections, we only considered hypotheses where at least two different QTLs in that cluster were associated with that GO term. We calculated $p$-values for GO term enrichment in a cluster using a one-sided hypergeometric test, and corrected for multiple hypothesis testing using the Benjamini-Hochberg method with an FDR/adjusted $p$-value threshold of 0.05. We found 90 significant hypotheses at the 0.05 significance threshold; these 90 significant hypotheses correspond to 89 different GO terms (one GO term is significantly enriched in two different clusters and 3 of the 89 GO terms correspond to the "CUSTOM" Ras/PKA/TOR pathway related GO terms we defined in Table S5) and 16 different clusters.

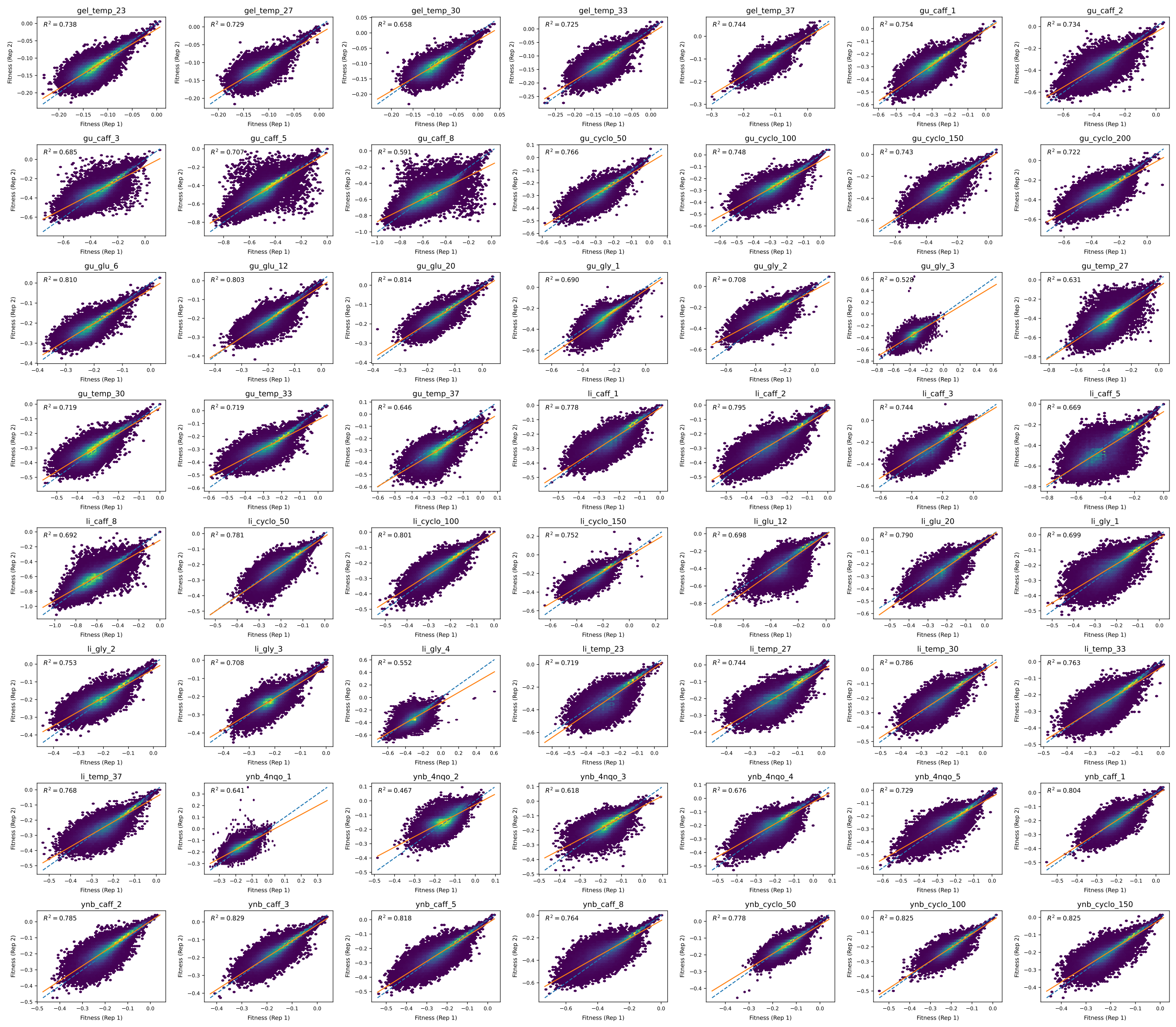


**Fig. S1. Accuracy of our fitness measurements**. Each plot shows inferred fitness of each segregant across two replicate phenotype assays in one of our 101 environments, along with 8 additional environments which were excluded due to poor quality data. The squared correlation coefficient ($R^2$), 1:1 line (dashed blue line), and best fit line (solid orange line) are also shown in each plot. Points on these hexbin plots are colored by density of fitness values.

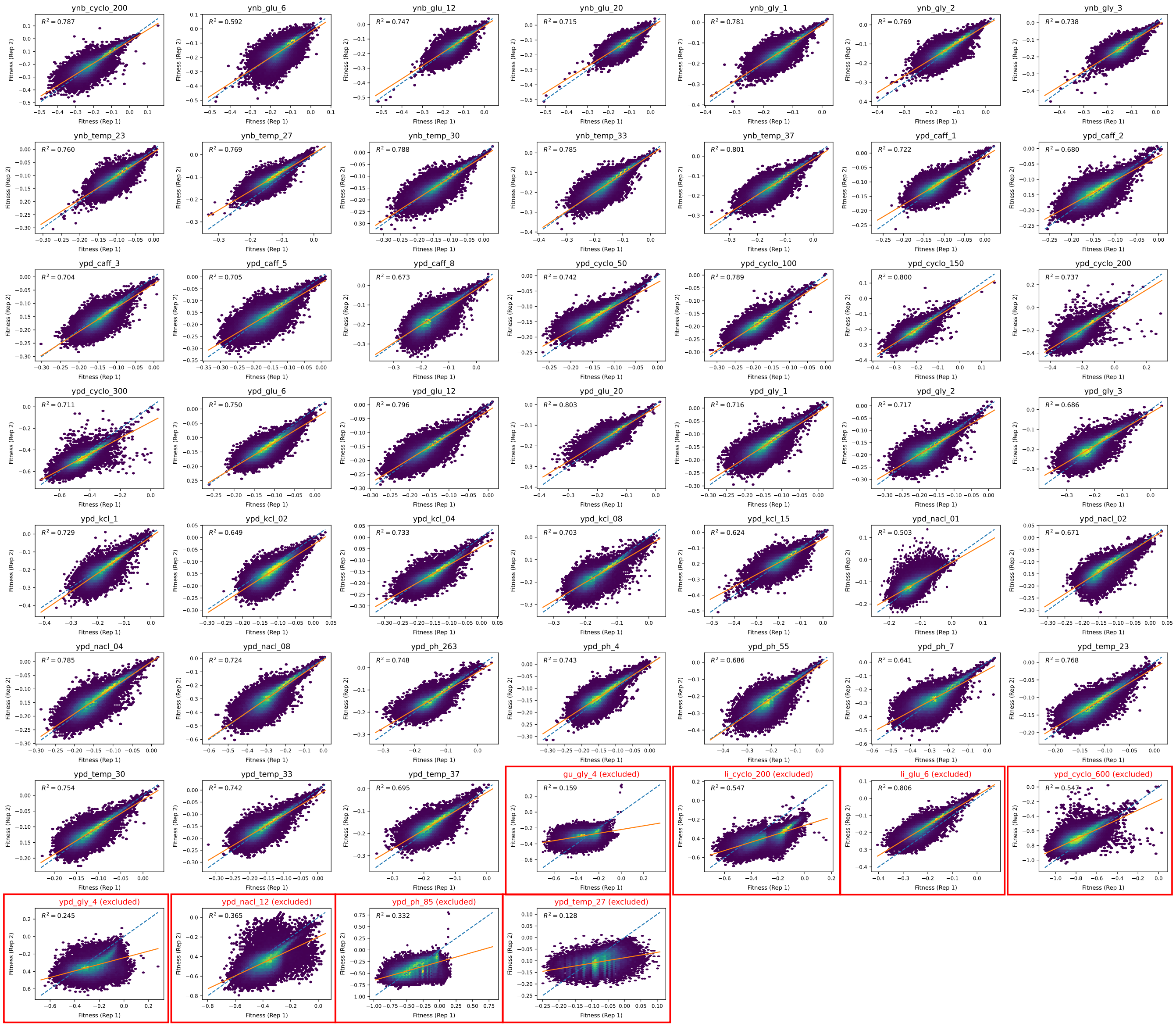


**Fig. S1.** [Continued]

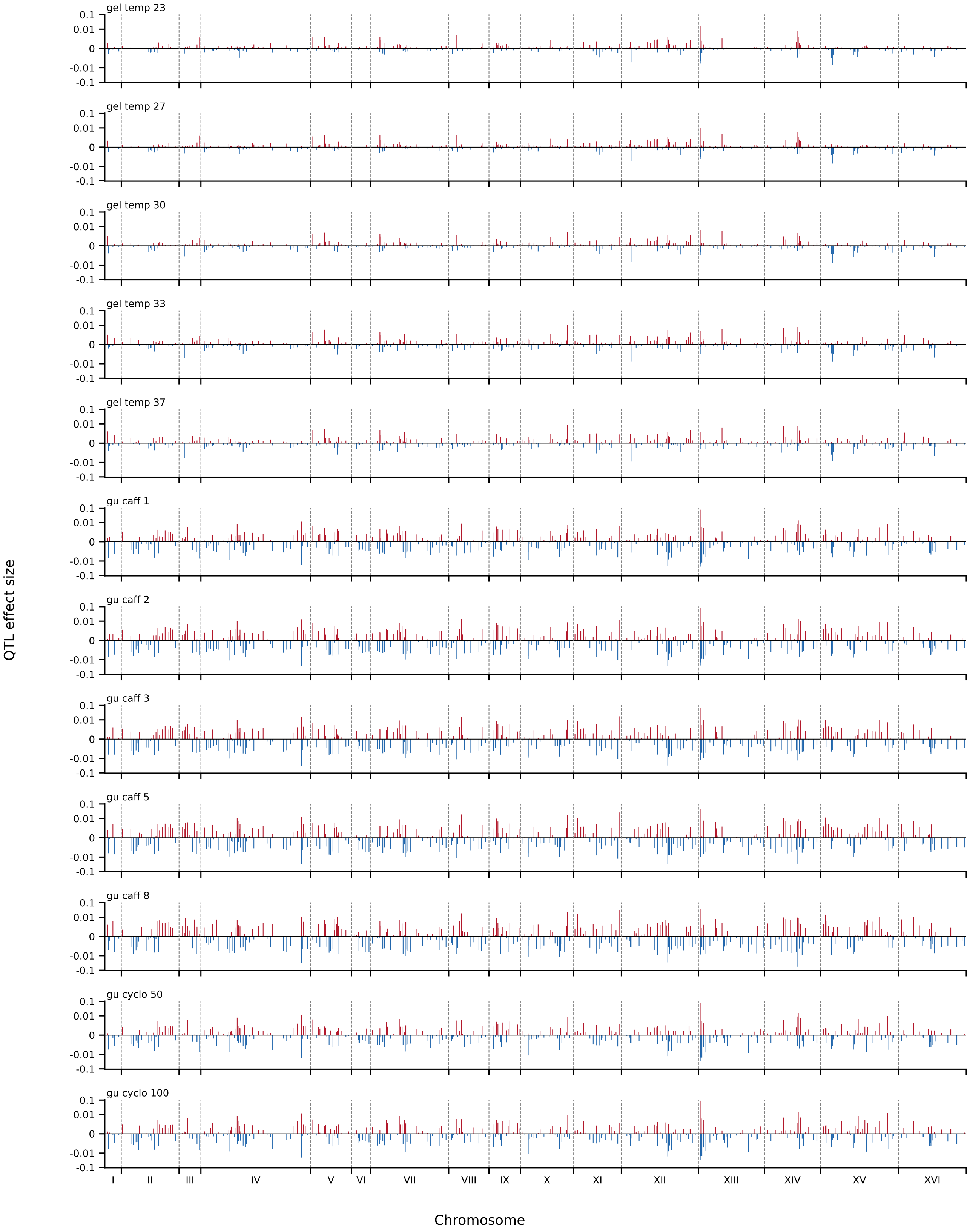


**Fig. S2. The genetic architecture of the measured growth rate phenotypes.** Each panel shows inferred positions of each QTL and their corresponding effect sizes in one of our 101 environments. Positive effect sizes (RM allele is more fit) are shown in red and negative effect sizes are shown in blue.

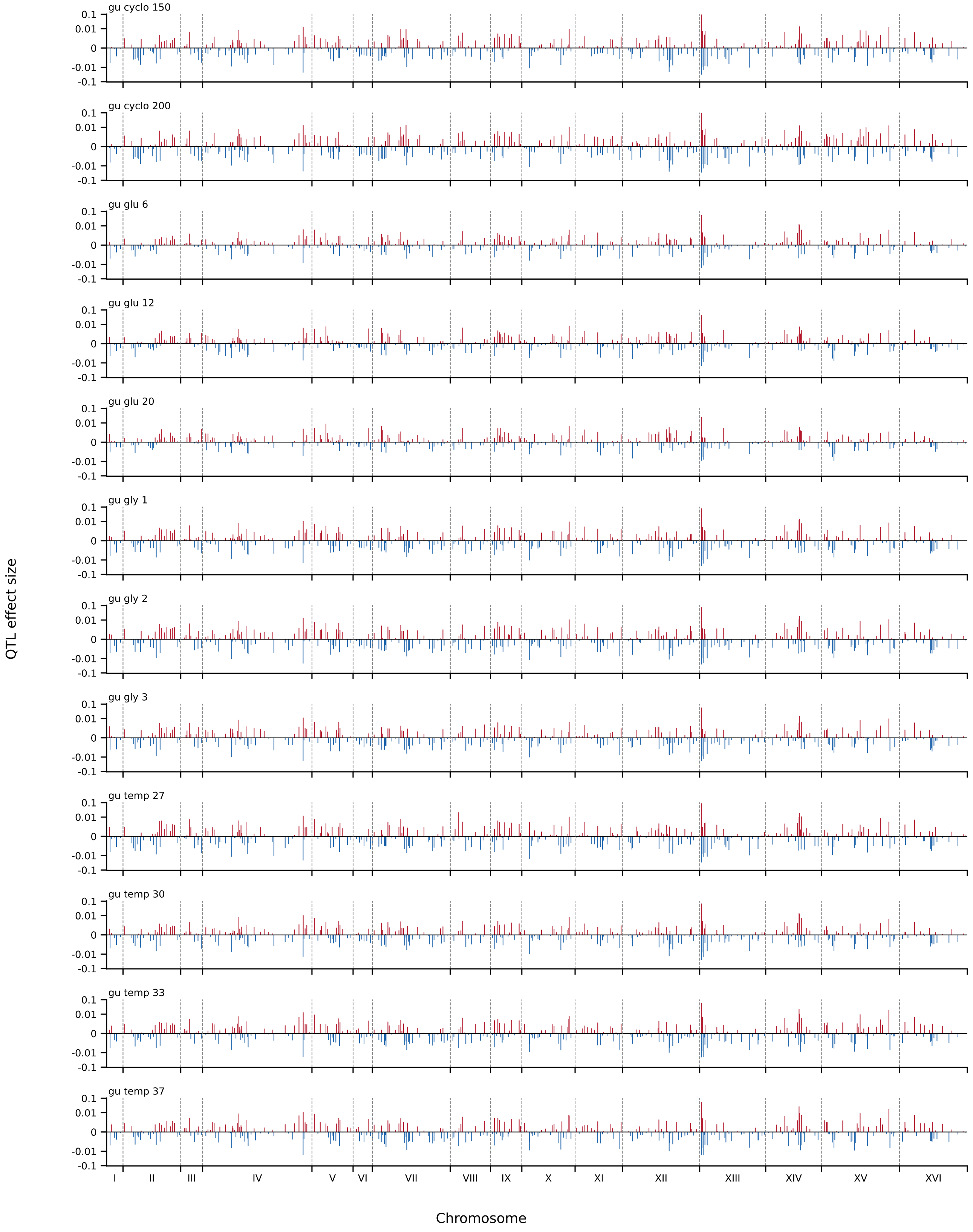


**Fig. S2.** [Continued]

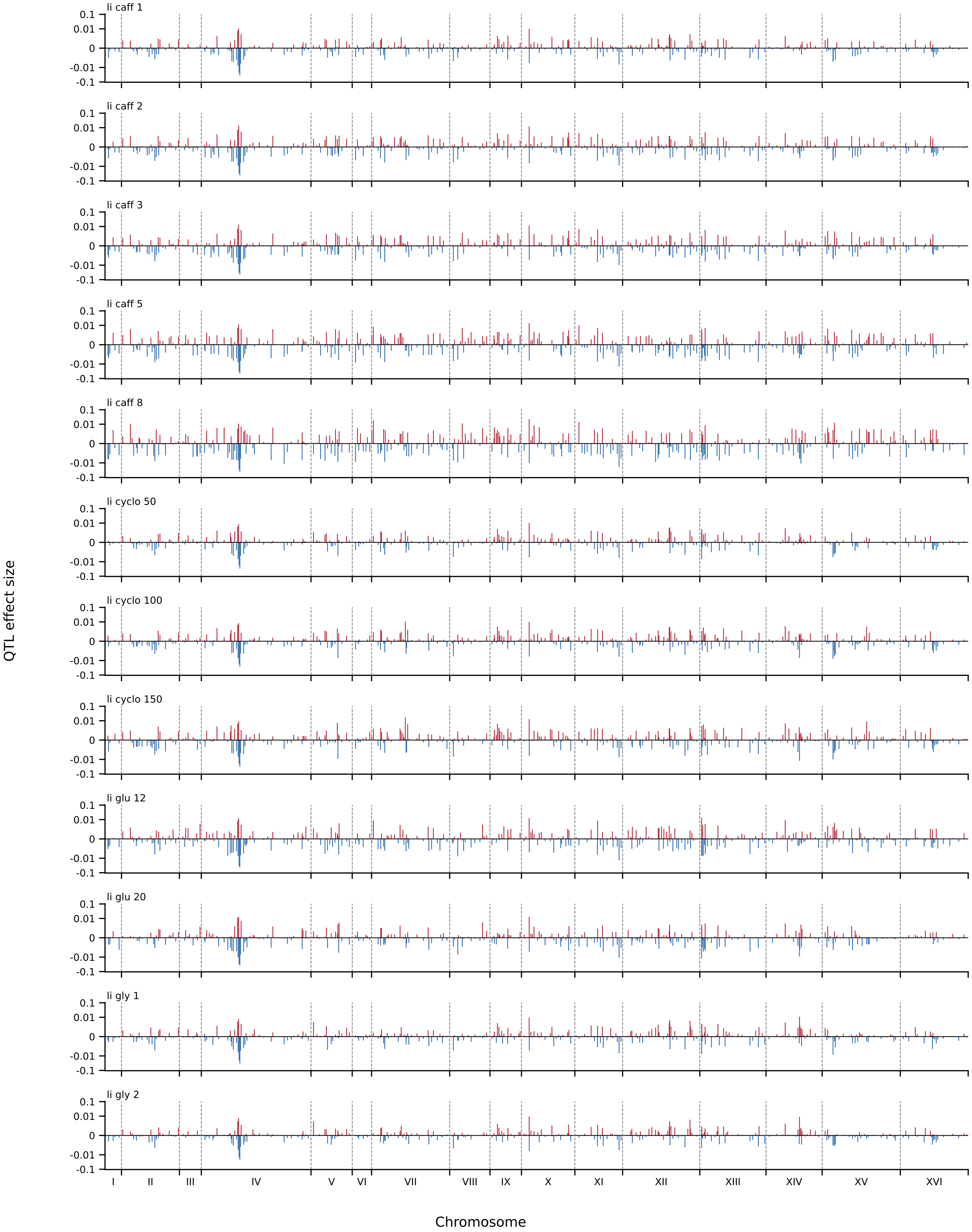


**Fig. S2.** [Continued]

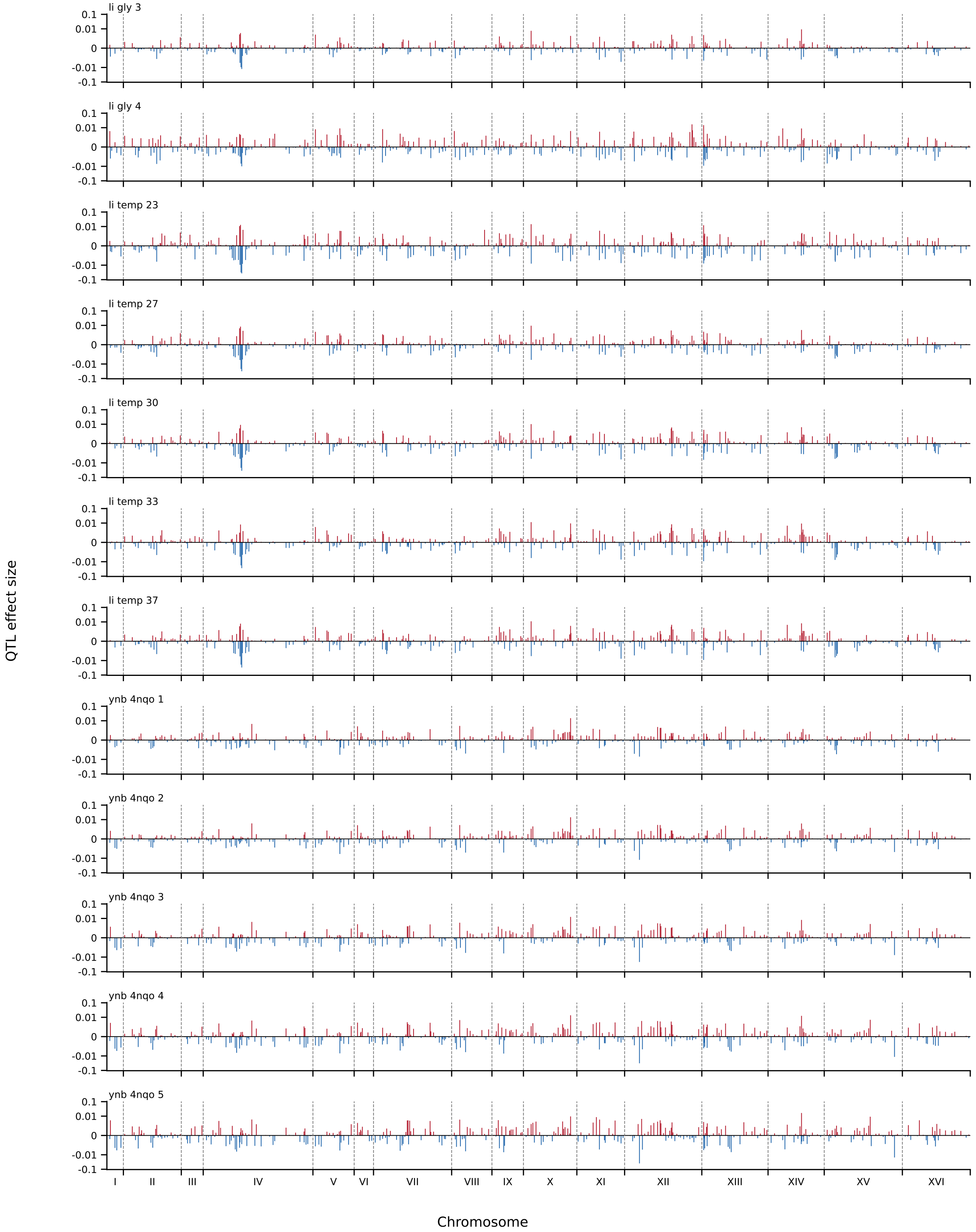


**Fig. S2.** [Continued]

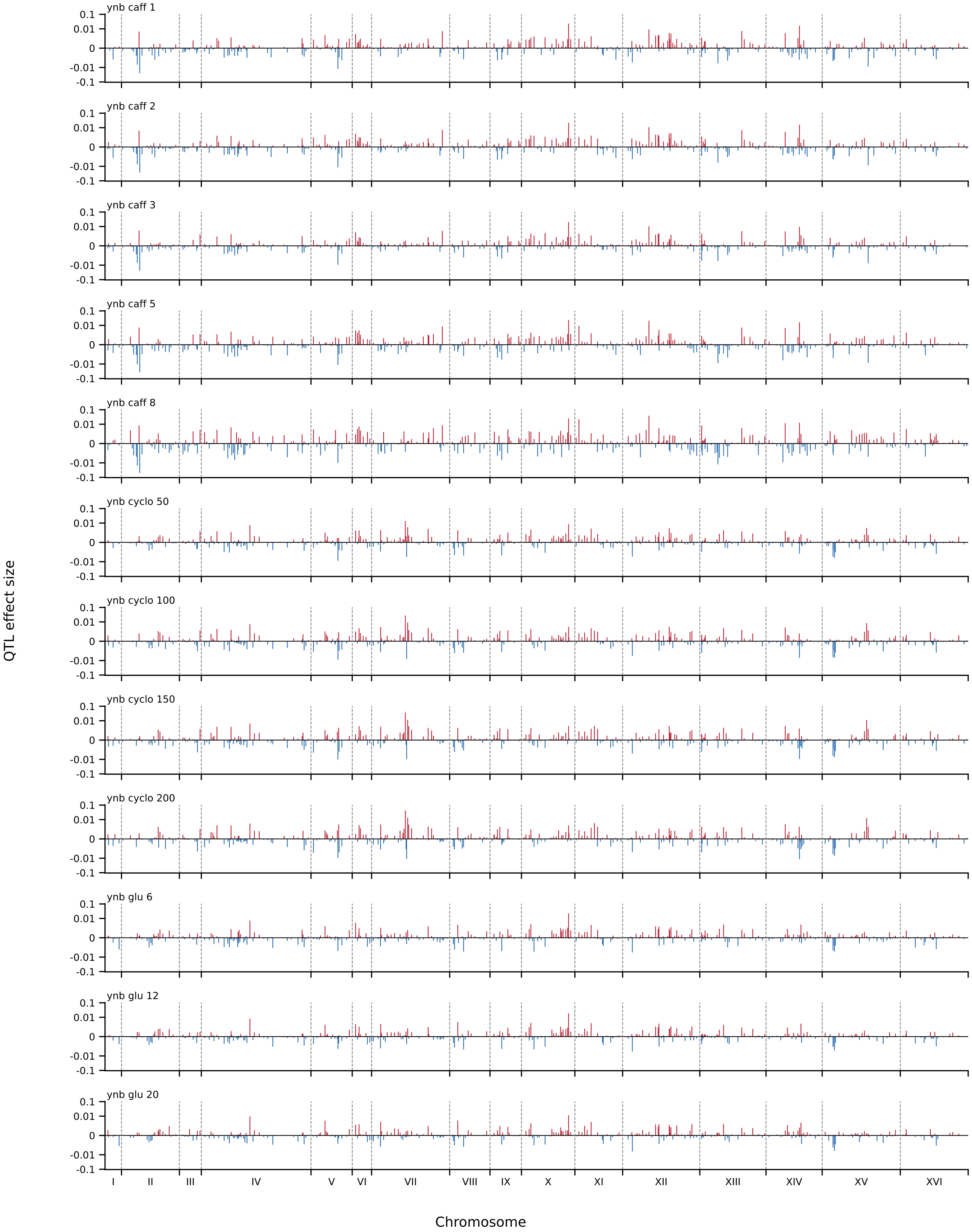


**Fig. S2.** [Continued]

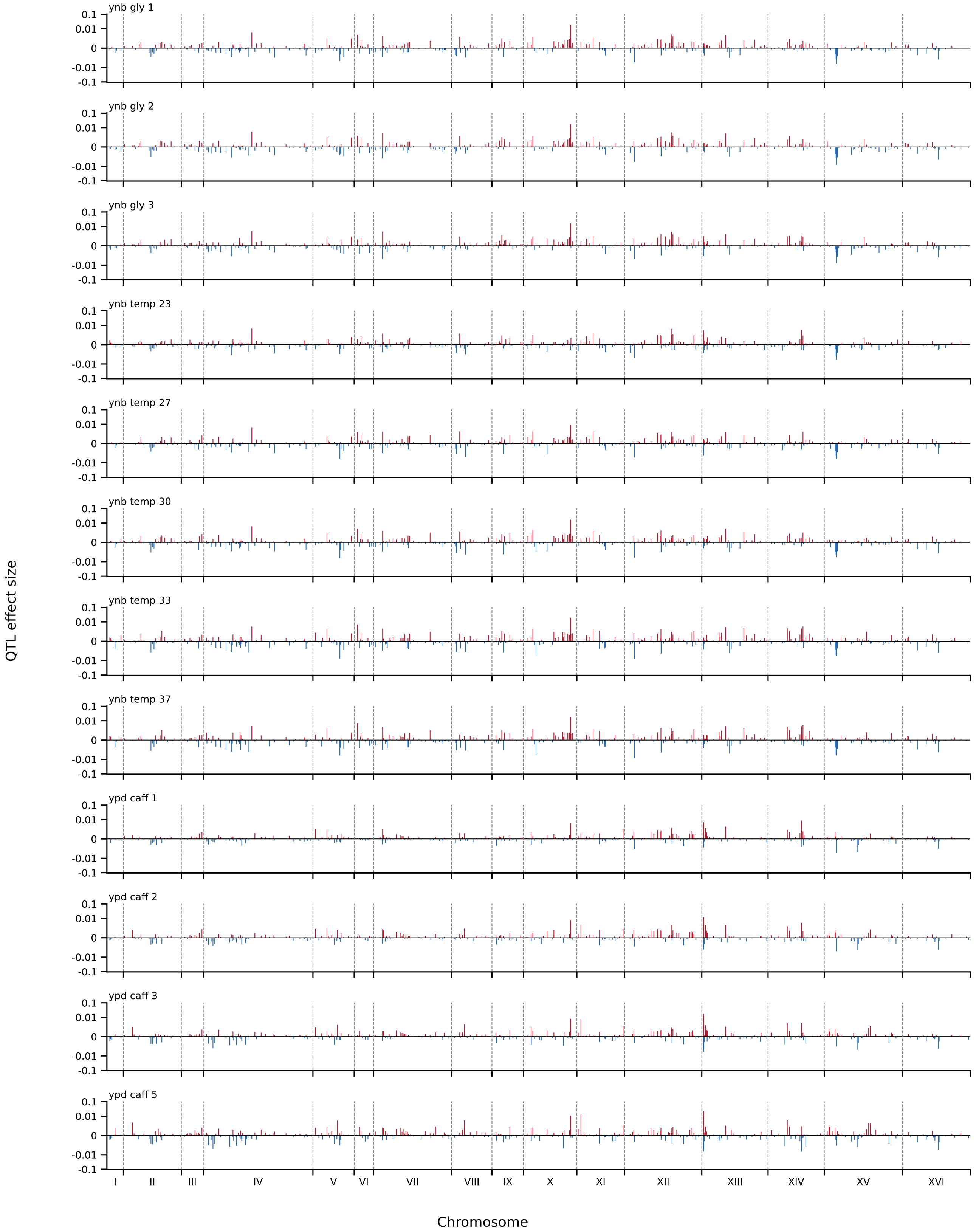


**Fig. S2.** [Continued]

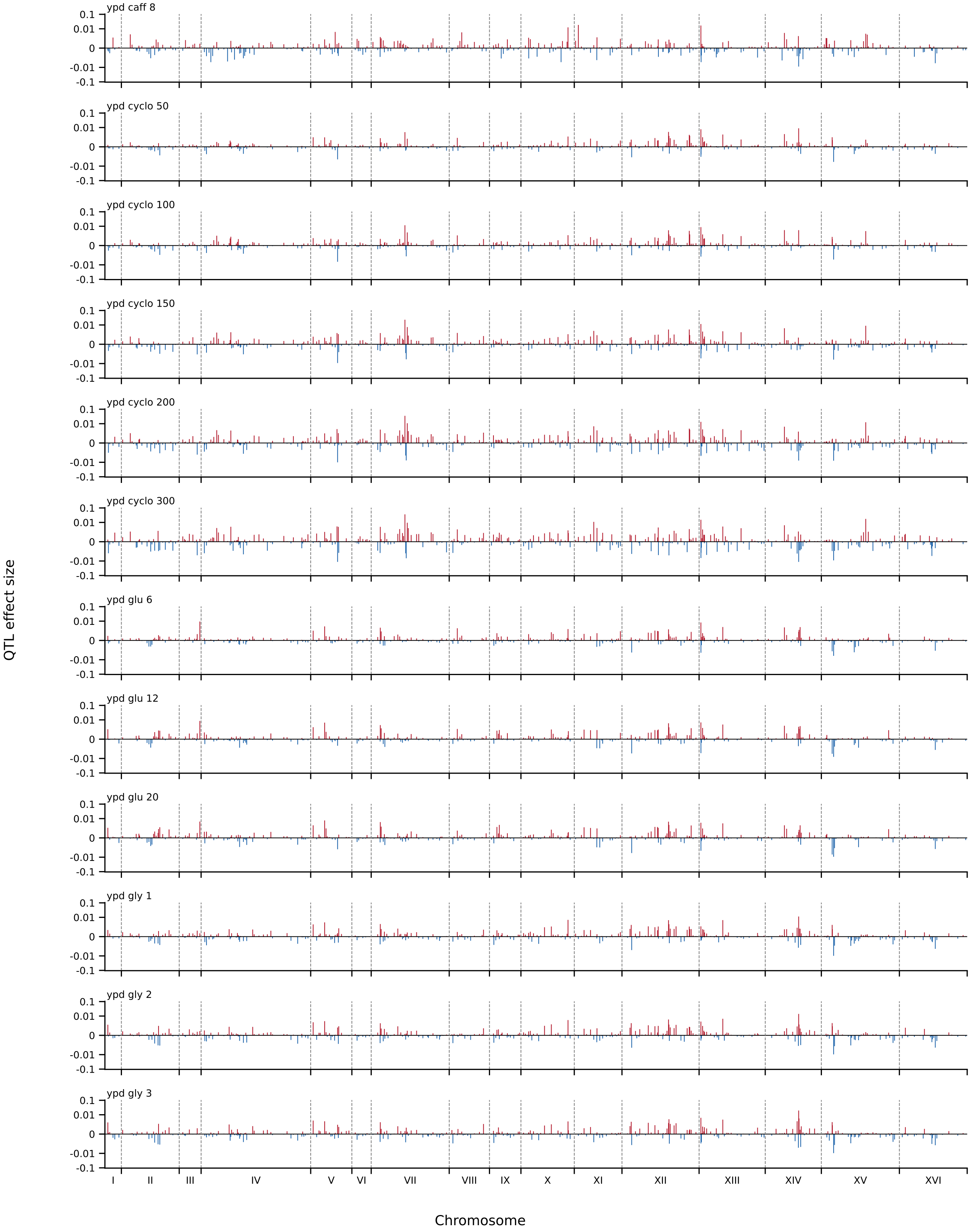


**Fig. S2.** [Continued]

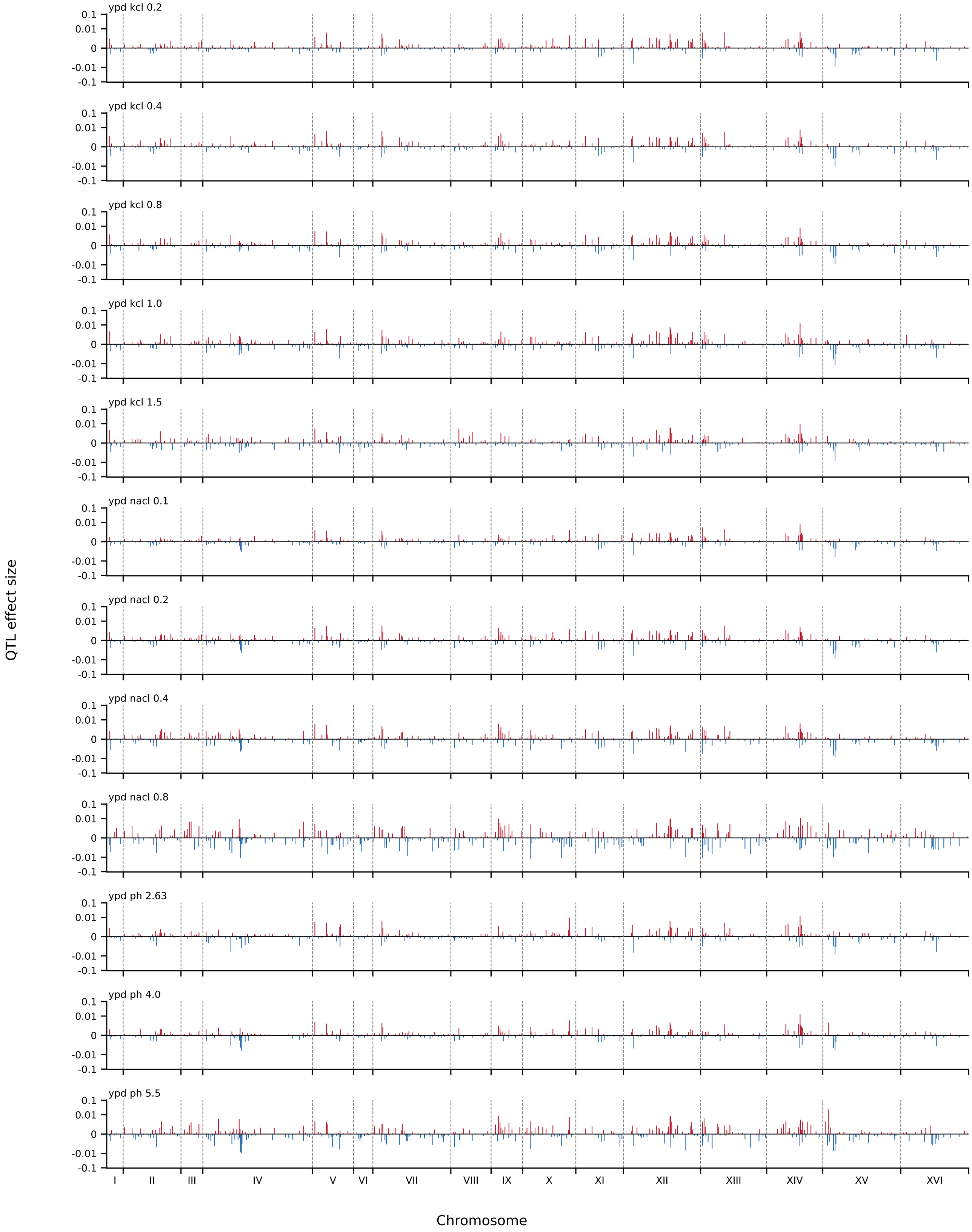


**Fig. S2.** [Continued]

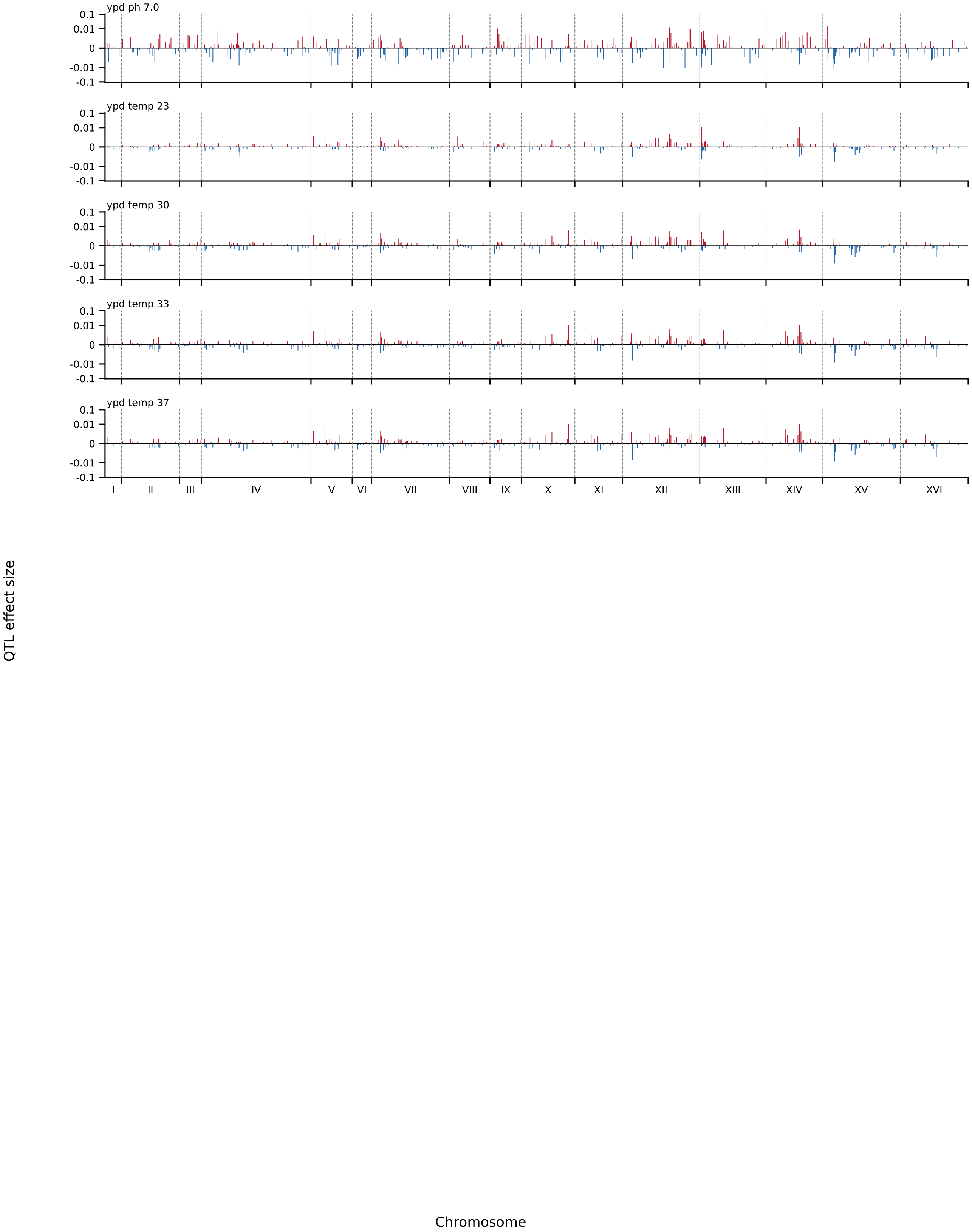


**Fig. S2.** [Continued]

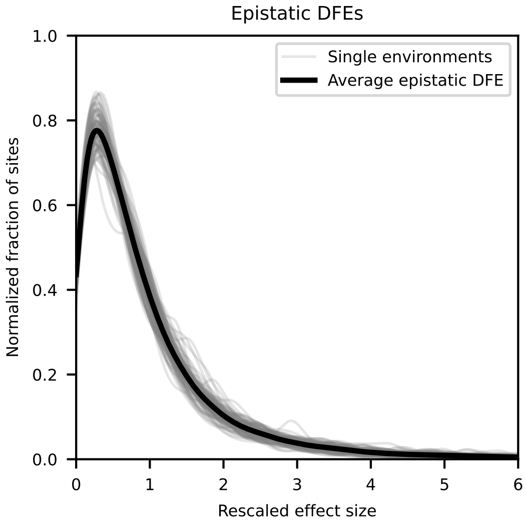


**Fig. S3. Distribution of pairwise epistatic effect sizes.** Each individual environment is shown in light gray (normalized by the mean effect size in that environment); average across all environments shown in black.

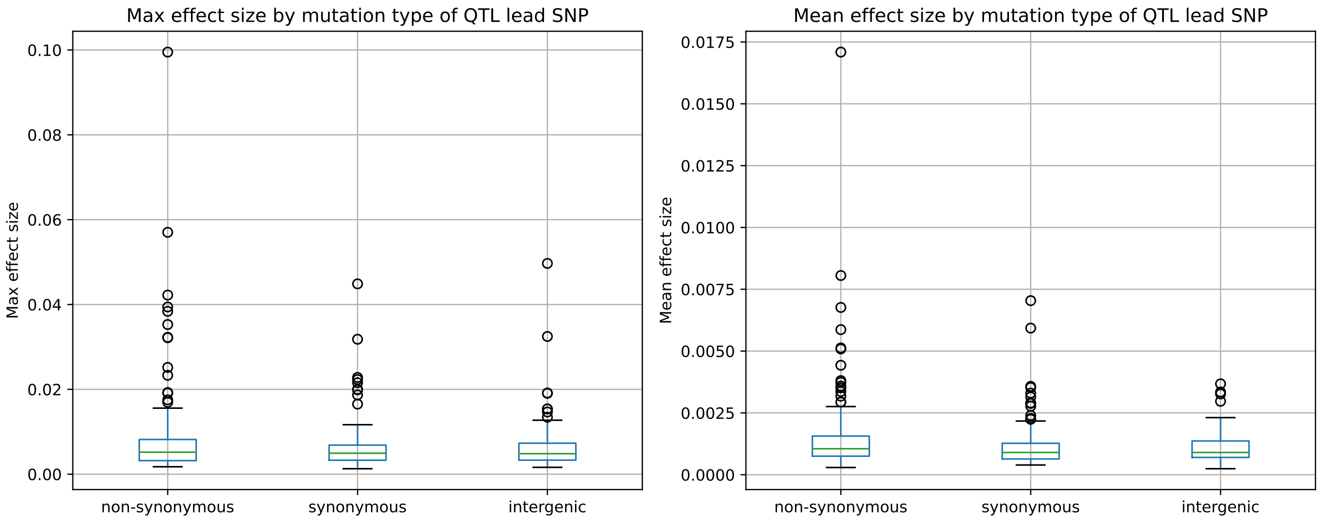


**Fig. S4. Effect sizes of QTLs classified by lead SNP mutation type**. Boxplots show the distribution of the maximum absolute effect size of a QTL across all 101 environments (left panel) and the mean absolute effect size of a QTL across all 101 environments (right panel), binned by the mutation type of the QTL lead SNP: non-synonymous, synonymous, and intergenic QTL lead SNPs.

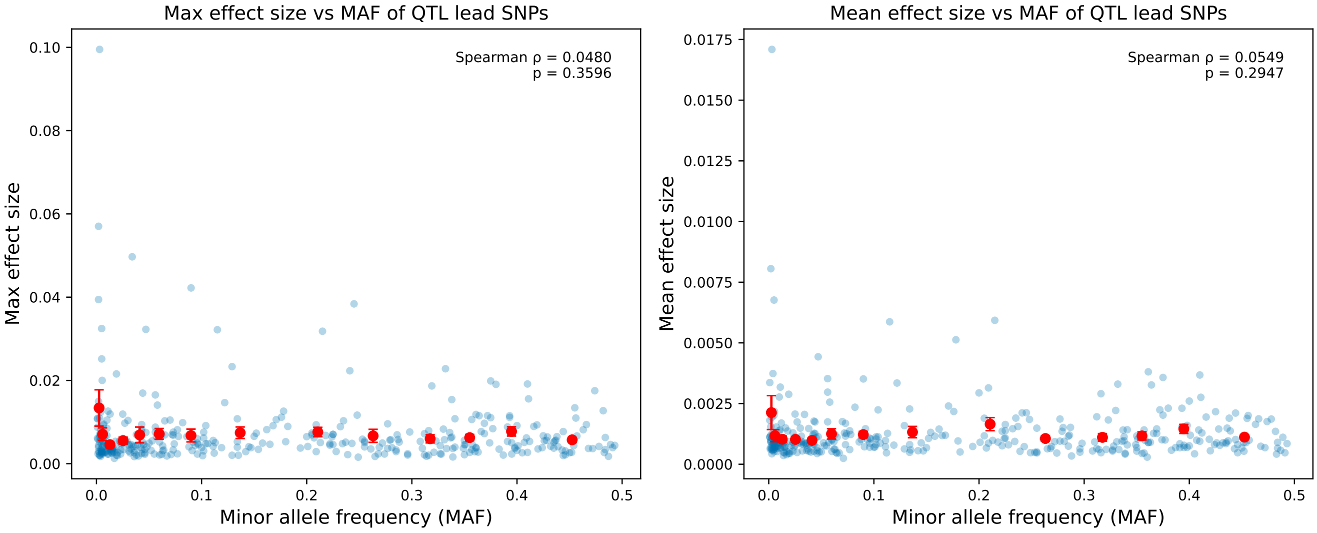


**Fig. S5. QTL effect sizes as a function of allele frequency**. Scatter plots show the maximum (left panel) and mean (right panel) QTL effect sizes across all environments plotted as a function of the lead SNP's minor allele frequency in the 1011 yeast genomes collection (blue points). The Spearman rank correlation $\rho$ and the $p$-value for this correlation are shown on the top-right corner of each plot. The red points show the mean y-axis value (+/- SEM) in bins of MAF (bins each contain 25 QTLs).

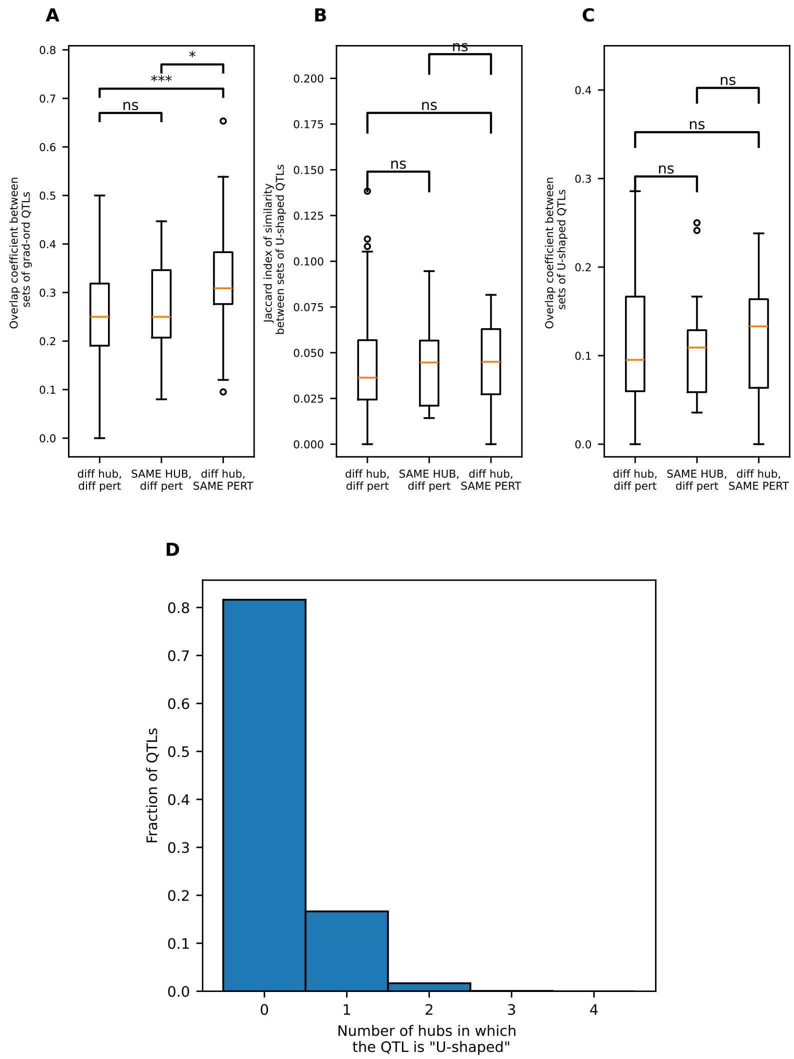


**Fig. S6. Similarity in the additive genetic architecture across the different perturbation gradients in our study. (A)** Overlap coefficient (a metric of similarity) between the sets of gradient-ordered QTLs shared between different types of pairs of gradients. Here * indicates $0.01 < p < 0.1$, ** indicates $0.001 < p < 0.01$, *** indicates $p < 0.001$, and ns indicates $p > 0.1$. **(B)** Jaccard index of similarity between the sets of U-shaped QTLs shared between the three different types of pairs of gradients. **(C)** Overlap coefficient of similarity between the sets of U-shaped QTLs shared between the three different types of pairs of gradients. **(D)** Histogram showing the fraction of QTLs that have U-shaped effects across 0, 1, 2, 3 or 4 different hub environments with the same perturbation gradient added.

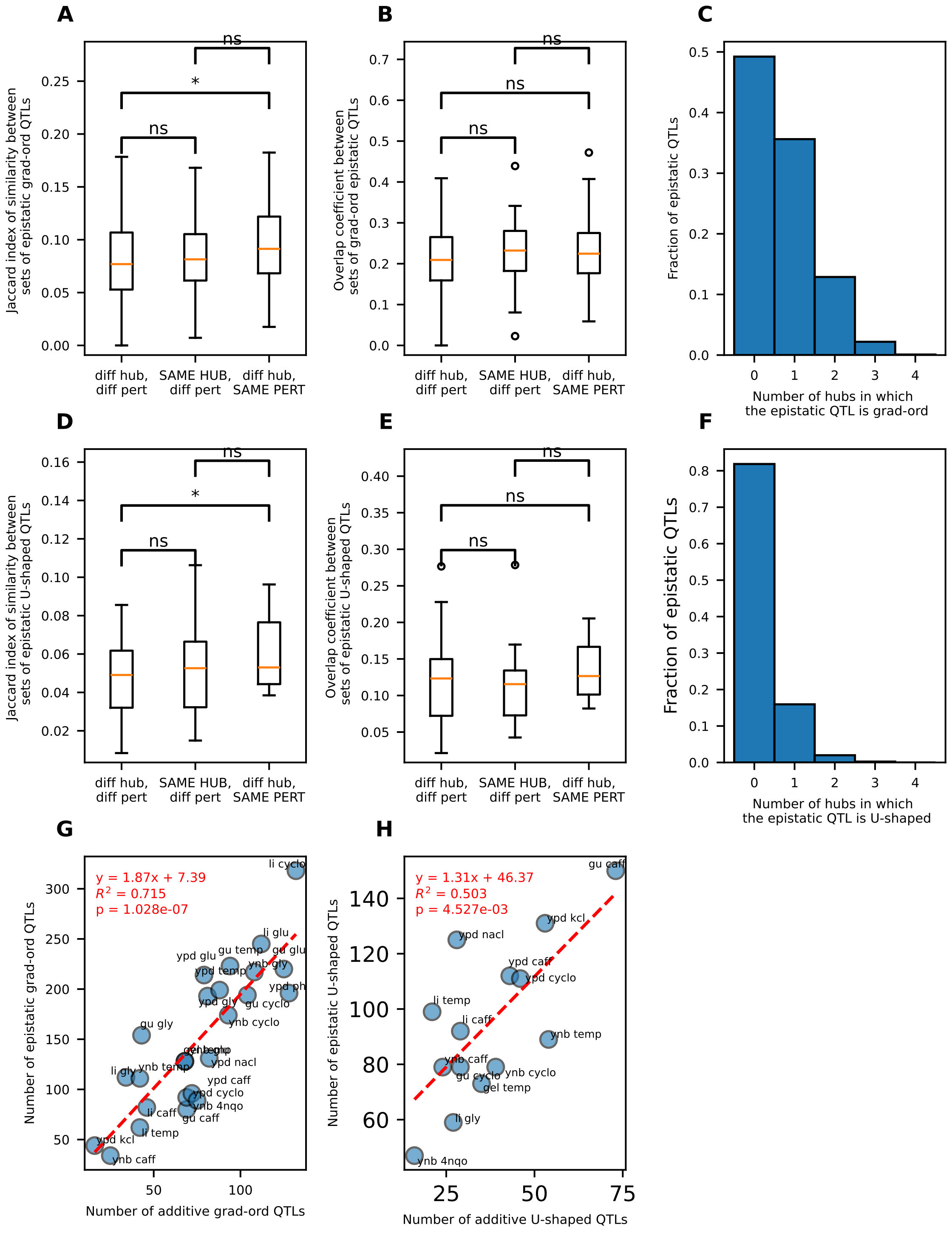


**Fig. S7. Similarity in the pairwise epistatic genetic architecture across the different perturbation gradients in our study. (A)** Jaccard index of similarity between the sets of gradient-ordered pairwise epistatic QTLs shared between the three different types of pairs of gradients. **(B)** Overlap coefficient of similarity between the sets of gradient-ordered pairwise epistatic QTLs shared between the three different types of pairs of gradients. **(C)** Histogram showing the fraction of pairwise epistatic QTLs that are gradient-ordered in 0, 1, 2, 3, or 4 different hub environments with the same perturbation gradient added. **(D)** Jaccard index of similarity between the sets of U-shaped pairwise epistatic QTLs that are shared between the three different types of pairs of gradients. **(E)** Overlap coefficient of similarity between the sets of U-shaped pairwise epistatic QTLs that are shared between the three different types of pairs of gradients. **(F)** Histogram showing the fraction of pairwise epistatic QTLs that have U-shaped effects in 0, 1, 2, 3, or 4 different hub environments with the same perturbation gradient added. **(G)** Number of pairwise epistatic gradient-ordered QTLs as a function of the number of additive gradient-ordered QTLs in each of the 25 different perturbation gradients in our study (the gradient name can be found next to each point in the plot). **(H)** Number of pairwise epistatic U-shaped QTLs as a function of the number of additive U-shaped QTLs in each of the 14 eligible perturbation gradients in our study. Note that the significance symbol convention used in this figure is the same as the one used in Fig. S6.

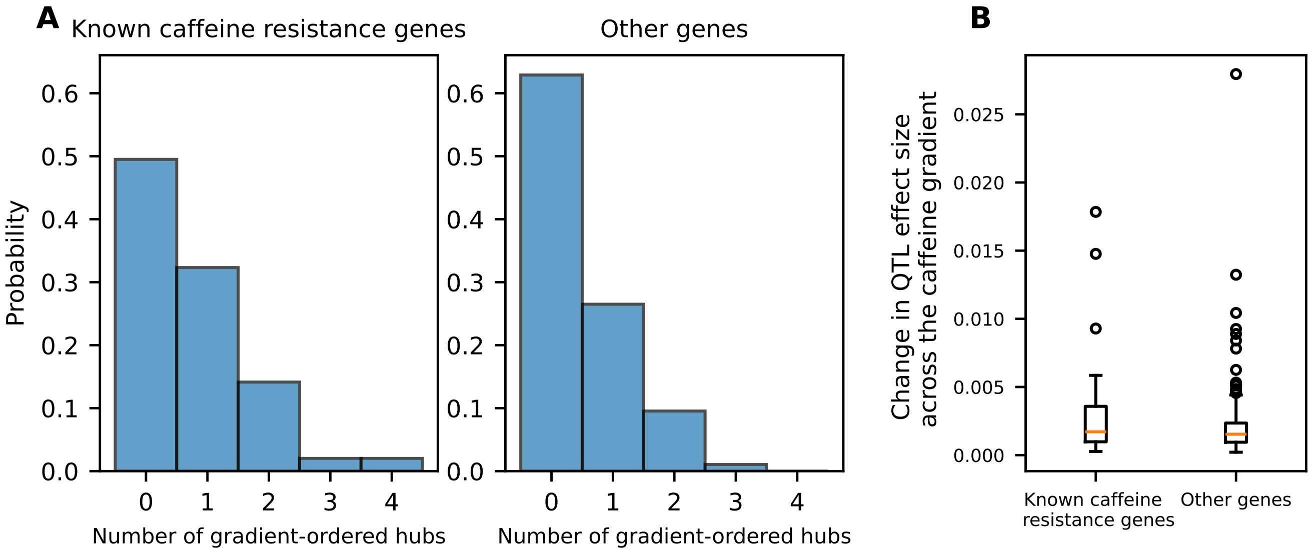


**Fig. S8. Gradient ordering across the caffeine gradient**. **(A)** Histogram showing the fraction of QTLs in known caffeine resistance genes (left) and in genes other than known caffeine resistance genes ("other genes"; right) that are gradient-ordered in the caffeine perturbation across 0, 1, 2, 3, or 4 different hub environments. **(B)** The distribution of change in QTL effect size across the caffeine gradient for QTLs located in known caffeine resistance genes vs QTLs located in other genes.

**Table S1 (TableS1.xlsx)**

List of environments in the order they are presented in Fig. 1D, along with details of the corresponding media and the pairwise fitness correlations (R) across all environments.

**Table S2 (TableS2.xlsx)**

Annotated list of all inferred QTLs and GO term enrichments and depletions.

**Table S3 (TableS3.xlsx)**

Summary of all gradient-ordered and U-shaped additive and epistatic QTLs in all of the eligible gradients.

**Table S4 (TableS4.xlsx)**

List of all additive QTLs annotated with whether their lead SNP is located in a known cycloheximide/4-NQO/caffeine resistance gene and the gradient-related quantities plotted in Fig. 4.

**Table S5 (TableS5.xlsx)**

Table of all z-scored effect size correlations between QTLs shown in Fig. 5, along with raw and adjusted p-values for all GO enrichment hypotheses tested for the clusters in this figure. Also contains a list of all additive QTLs annotated with tier 3 GO terms, and a list of Ras/PKA and TOR pathway subcomponents and the associated genes.

**Dataset S1 (DatasetS1_jointlyinferredfitnesses.tsv.gz)**

Jointly inferred fitness (across two replicate fitness assays) for all 99,950 F1 segregants in the 101 different environments.

**Dataset S2 (DatasetS2_replicatefitnesses.tsv.gz)**

Fitness inferred in each replicate fitness assay for all 99,950 segregants in the 109 different environments shown in Fig. S1.

**Dataset S3 (DatasetS3_all_SNPs_annotated.tsv.gz)**

Annotated list of all SNPs segregating in our BY x RM cross.
